# Metal toxicity contributes to the structuring of bacterial communities in the Arabidopsis phyllosphere

**DOI:** 10.64898/2026.04.12.717999

**Authors:** Jan F. Plewka-Mandelkow, Anna S. Thomas, Julia A. Vorholt, Ute Krämer

## Abstract

- The causal factors shaping plant-associated microbiota are incompletely known. Elevated concentrations of the micronutrients zinc (Zn), manganese (Mn) and copper (Cu), and exposure to non-essential trace elements including cadmium (Cd) and arsenic (As), can be toxic. Here we explored whether differences in metal(loid) sensitivity between plants and bacteria influence phyllosphere bacterial community composition.
- 224 representative *Arabidopsis thaliana* phyllosphere bacterial strains were screened on metal(loid) concentration series in synthetic media. We obtained leaf apoplastic fluid ionomes for comparisons with bacteriotoxicity profiles, and tested for relationships between strain-wise metal(loid) tolerances, phylogeny and gene content.
- Leaf apoplastic Zn^2+^ and Cd^2+^ concentrations were the most likely to arrest growth of metal-sensitive bacteria *in planta*. Soil bacterial strains were several-fold more sensitive to both these metals than leaf strains, consistent with selection for increased bacterial Zn and Cd tolerance in the phyllosphere. Strains known to govern bacterial community structure were metal-sensitive, with only minor influences of between-metal and between-strain interactions. Bacterial genus explained considerable proportions of the variances in metal(loid)-related gene content and tolerance phenotypes. Bacterial Cd tolerance correlated with the presence and copy number of known Cd-related genes.
- Our results suggest that plant metal homeostasis contributes to structuring bacterial communities in the leaf endosphere.

## Introduction

Diversity among bacteria is unparalleled, and they have colonized the broadest range of habitats in the world, including soil, water, and multicellular organisms. The plant-associated microbiota comprises complex microbial communities influenced by the plant host, microbe-microbe interactions, and other environmental factors, and they can have positive effects on plant fitness (Müller *et al*., 2016b; Ritpitakphong *et al*., 2016; Vogel *et al*., 2021). The phyllosphere is one of the largest habitats for bacteria with an estimated global area of around 1 billion km^2^ (Vorholt, 2012) and comprises two distinct microbial habitats: one on the outer leaf surfaces named the phylloplane, and one inside the leaf named the endosphere. The leaf endosphere is accessible from the leaf surface *via* the stomata while they are open. Within a given plant organ, the volume outside the plasma membranes bordering all cells is referred to as the apoplast, comprising both the cell walls and the intercellular space in between (Sattelmacher, 2001). In general, the leaf apoplast is at least partially filled with an extracellular solution, and the remainder of it is occupied by air. In addition to its role as a habitat for endosphere microbial communities, the apoplast serves as a site for the inter-cellular movement of gases, water and solutes, as well as for the storage and detoxification of organic and inorganic compounds including metals (Küpper *et al*., 2000; Sattelmacher, 2001; Kajala *et al*., 2019).

Impacts on bacteria inhabiting the phylloplane, rhizosphere and rhizoplane are determined predominantly by the environment around the plant. The closer the plant host, the stronger is its influence on the microbiota (Bulgarelli *et al*., 2013; Chen *et al*., 2020). The plant begins to exert some control over the composition of its apoplast progressively from the root epidermis inwards, where both diffusion from the soil solution and microbial colonization can occur easily on the external side of apoplastic barriers, or during root developmental stages prior to their formation. In roots, apoplastic barriers are constituted by the Casparian strip that is positioned radially around root endodermal cells as well as the suberization of the outwardly oriented surfaces of epidermal and endodermal cell layers, both initiated at specific stages of differentiation. Endodermal suberization is plastic to some degree and regulated dependent on both the plant’s nutritional status and plant-associated microbial communities (Robb *et al*., 1991; Andersen *et al*., 2015; Barberon *et al*., 2016). The apoplast on the internal side of the root apoplastic barriers and of the leaf endosphere represent comparably sheltered niches for the inhabiting microbes. At these sites, the plant host has a predominant influence on apoplast composition, thus providing opportunities for niche construction by the plant. To date, little is known about the composition of this micro-habitat and any evolutionary adaptations to it in the inhabiting microbes.

The plant apoplast is known to function in the storage and detoxification of excess metal cations, including those of so-called heavy metals, or – more accurately – class B (e.g., Cu^+^) and borderline elements (e.g., Cu^2+^, Cd^2+^, Fe^2+^, Fe^3+^, Mn^2+^, Zn^2+^ (Nieboer, 1980; Farvardin *et al*., 2020)). Among these, the micronutrients (mostly Fe, Zn, Cu, Mn and Ni) are required in small quantities by all organisms to serve as or within protein cofactors acting in ligand binding and in the catalysis of a variety of chemical reactions (Pais & Jones, 1997; Marschner & Marschner, 2012). Because of their powerful chemical properties, these same nutrients can become highly toxic even when present in slight excess. Moreover, chemically similar non-essential trace elements are ubiquitously present in the environment, for example Cd. All organisms, including bacteria and plants, possess metal homeostasis networks which operate to fulfill nutritional requirements while counteracting metal toxicities across their specific environmental ranges (Clemens, 2001; Nies & Grass, 2009; Chandrangsu *et al*., 2017). Over the course of evolution, metal homeostasis networks, as well as nutritional metal needs, have diverged between bacteria, fungi and plants to some extent. For example, the metalloproteome of plants generally requires more Cu and Zn, and less Ni, than that of most bacteria (Williams, 2007). This divergence implies that metal concentrations maintained and tolerated by the plant within the apoplast may be inhibitory to bacterial colonists, raising the possibility that plants modulate apoplastic metal levels to selectively influence microbial community composition. In support of this, *Arabidopsis thaliana* Zn^2+^ efflux ATPases HMA2 and HMA4 are required for resistance to the necrotrophic fungal pathogen *Plectosphaerella cucumerina* (Escudero *et al*., 2022), for example, demonstrating that plant-controlled apoplastic metal levels can shape interactions with invading microbes. Moreover, high leaf apoplastic zinc concentrations can directly protect the metal hyperaccumulator *Noccaea caerulescens* (ecotype Prayon) against infection by *Pseudomonas syringae* pv. maulicola M4 (Fones *et al*., 2010). The concept of nutritional immunity is well-established through medical research, whereby the host manipulates nutrient availability to limit pathogen growth. However, related concepts are less well understood and remain underexplored in plant science (Fones & Preston, 2013; Murdoch & Skaar, 2022; Krämer, 2024).

Here we explored the relationship between metal(loid) levels in the plant apoplast and plant-associated microbial communities. We profiled the metal(loid) tolerances of 224 representative phyllosphere bacterial strains of the *At*-LSPHERE collection, a curated set of isolates from the model plant *A. thaliana* (Bai *et al*., 2015), focusing on elements that could reach bacteriotoxic levels inside leaves of plants in their natural non-contaminated environments: Cd, Cu, Mn, Zn, and arsenic. We then generated ionomic data for the Arabidopsis leaf apoplast, which we evaluated in the context of phyllosphere bacterial metal(loid) tolerances as well as in comparison to tolerance profiles of soil bacteria, and we identified genomic predictors of metal tolerance in phyllosphere bacterial strains. Combining these datasets, we assessed whether metal(loid) co-tolerance patterns reflect shared chemical properties or environmental selection. In summary, our work identifies the modulation of Zn and Cd levels in the leaf apoplast as a potential means for plants to shape the bacterial community in the leaf endosphere.

## Materials and Methods

### Biological materials and cultivation conditions

*Arabidopsis thaliana* (L.) Heynh., accession Columbia (Col-0), wild-type seeds (Lehle Seeds, Round Rock, TX, USA) were sown onto moistened standard soil (Type Minitray, Balster Einheitserdewerk, Fröndenberg, Germany) and stratified in darkness at 4°C for 2 d, followed by cultivation in 8-h light (145 μmol m^−2^ s^−1^, 22°C)/16-h dark (18°C) cycles in a growth chamber (GroBank BB-XXL.3, CLF Plant Climatics, Wertingen, Germany). After 2 w, single seedlings were transferred into 0.7-L pots containing standard soil or a mix of a natural soil from Bad Gottleuba, 50°50’45.7” N 13°59’43.1” E, with sand and standard soil (35 + 15 + 50 volume parts, respectively), and cultivated for another 5 to 6 w.

The 224 isolates of the *A. thaliana* phyllosphere (*At*-LSPHERE; Bai et al. 2015) collection and the 33 bacterial isolates from unplanted soil (Bai *et al*., 2015) were cultivated in 3 mL liquid R2A medium containing methanol (0.5% v/v) in darkness at RT, with continuous shaking (300 rpm), for 2 to 3 d. Saturated cultures were mixed 1:1 (v:v) with sterile-filtered 70% (v/v) glycerol in ultrapure water and stored at -80°C. All cultivation was conducted in R2A media containing methanol (0.5% v/v), prepared and autoclaved as described (Bai *et al*., 2015), and subsequently supplemented with metal(loid)s from sterile-filtered (Filtropur S 0.2 µm, SARSTEDT AG & Co. KG, Nürmbrecht, Germany) stock solutions where needed (Tables S1 and S2).

### High-throughput metal(loid) tolerance screening

Bacterial stocks were defrosted and screened on agar-solidified media in square polystyrene petri plates (120 x 120 x 17 mm) using a sterilized 96-pin microplate replicator (Boekel Scientific, Feasterville, USA), followed by incubation at RT in the dark. Growth was scored based on photographs taken after 10 d and 20 d (occasionally) of cultivation. Three independent experiments (repeats) were conducted, each comprising a control of unamended medium and a series of six concentrations for each metal(loid) (Table S2). Additional metal(loid) concentrations were tested in further experiments.

### Quantification of Metal(loid) Tolerances in Single Strains and Synthetic Communities

Single colonies of keystone strains (Carlström *et al*., 2019) and of each strains of the synthetic communities (SynComs) were transferred into 3 mL of liquid medium and pre-cultivated at RT for 1 to 2 d. Each culture was diluted with sterile liquid medium to an OD_600_ of 0.2 (SYNERGY HTX multi-mode reader, BioTek Instruments GmbH, Bad Friedrichshall, Germany), mixed 1:1 (v:v) with 70% (v/v) sterile-filtered glycerol, with equal volumes used per strain to assemble each of the SynComs, and stored at -80°C. For the quantification of metal tolerances based on growth curves, five µL of each stock were added per well containing 195 µL liquid medium with or without (control) metal supplementation in a 96-well plate. Bacterial strains were cultivated in darkness at 22 to 28°C for ≥ 20 h (Cd tolerance) and 48 h (combined metal toxicities), with one OD_600_ measurement per h (5 technical replicates per experiment, 3 independent experiments) or per 0.5 h (SynComs; 6 technical replicates per experiment, 5 independent experiments) preceded by horizontal shaking at 200 rpm for 5 min.

### Isolation of leaf apoplastic wash fluid from *A. thaliana* and Multi-Element Analyses

Apoplastic wash fluid was prepared as described (O’Leary *et al*., 2014), with small modifications. Freshly harvested leaves were immersed in an infiltration fluid (IF) of either ultrapure water or 100 µM EDTA (pH 7.2), followed by three cycles of vacuum-infiltration at 30 mbar for 3 min and release of the vacuum over a period of 3 min, with recovery of the AWF by centrifugation in a swinging bucket rotor at 1,000x*g* and 4°C for 15 min. We experimentally determined a dilution factor for the AF in the AWF (2.35; *n* = 4) (O’Leary *et al*., 2014). For multi-element analyses, dry aliquots of AWF, leaf powder, liquid media, agar and soils were acid-digested, followed by Inductively Coupled Plasma-Optical Emission Spectrometry (ICP-OES) using an iCAP Duo 6500 (Thermo Fisher, Dreieich, Germany) (Stein *et al*., 2017) (see also Supporting Methods).

### Data Analyses and Visualization

Data were organized in Microsoft Excel (Microsoft Corporation, Redmond, USA), with analysis using R (version 4.3.0) (R_Core_Team, 2020) in the RStudio environment (version 2023.06.0.421) (RStudio_Team, 2021) and plot post-processing in Inkscape (version 1.2.2) (Inkscape_Project, 2020). Dose-response curves (*f*(*x*) = *E_inf_+(1-E_inf_)/(1+(x/EC_50_)^h^)*) and metrics (*EC_n_* = *EC_50_ × [(0.01 n)/(1 – 0.01 n – E_inf_)]^1/h^*) were obtained using the `GRfit()` function of the *GRmetrics* package (*E_inf_* : proportion of bacteria surviving an infinite metal concentration, set to 0 for calculating EC*_n_*; EC*_n_*: metal(loid) concentration at which *n*% of the strains were unable to grow; *h*: Hill coefficient of the fitted curve; *x* the metal concentration in the medium [mM]; and *f*(*x*) the proportion of surviving strains). We calculated strain-wise MPC for each metal(loid) as the median of three independent experiments, followed by conversion to *z*-scores across all strains for each metal(loid) using `scale()`. A Euclidean distance matrix for hierarchical clustering was generated with the `dist()` function. Based on an agglomerative coefficient of 0.98 calculated using the `agnes()` of *cluster*, Ward clustering was the best-fitting method and implemented through `hclust()` using the ward.D2 method. Principal component analyses (PCAs) were conducted using `prcomp()`, with significance testing using *PCAtest*. Null distributions for PCA statistics were generated using 999 random permutations, and sampling variance was estimated based on 999 bootstrap replicates. The `capscale()` function of *vegan* was used for constrained principal coordinate analyses (CPCoA). Statistical significance of the ordinations and confidence intervals for the variance were calculated by ANOVA-like permutation tests using `permutest()` and `anova.cca()` (Robbins *et al*., 2018). We calculated growth rate *r* = *ln(OD_600_(T_2_)/OD_600_(T_1_))/(T_2_-T_1_)* and apparent doubling time *t_d_* = ln(2)/*r* from the first Log phase of growth between Time points T_1_ and T_2_ identified using the *zoo* package and a linear model (ln-transformed OD_600_). The phylogenetic tree was built with `bionj()`from *ape* using 16s rRNA sequences (Bai *et al*., 2015) (www.at-sphere.com) aligned with `msa()`followed by the computation of a distance matrix using `msaConvert()` from *msa* and `dist.alignment()` from *seqinr*.

## Results

### Profiles of metal(loid) tolerances among Arabidopsis phyllosphere bacterial strains

To obtain insights into whether the metal(loid) levels in the leaf apoplast can influence the composition of phyllosphere microbial communities, we profiled the tolerance limits of all individual strains within the publicly available representative collection of Arabidopsis leaf phyllosphere bacteria named *At*-LSPHERE (*n* = 224) (Bai *et al*., 2015). For comparison, we additionally screened a collection of bacteria from unplanted soil (*n* = 33) (Bai *et al*., 2015) in the same manner. We spotted bacterial suspensions of each strain on agar-solidified R2A media supplemented with concentration series of the essential metals copper (Cu^II^), zinc (Zn^II^), and manganese (Mn^II^), as well as of the non-essential trace metal cadmium (Cd^II^), and the metalloid arsenic (As) in two common inorganic forms, arsenite (As^III^) and arsenate (As^V^) (Tables S1 to S3).

Of all tested metal(loid)s, Cd^II^ was toxic for some *At*-LSPHERE strains at the lowest concentration (0.8 µM; Dataset S1). A concentration of 49 µM Cd eliminated the growth of 50% of the bacterial strains (termed here Effective Concentration EC_50_), with a factor of 130 between the Cd concentrations eliminating the 5% most sensitive (EC_5_) and the 5% most tolerant strains (EC_95_), defined here as concentration range (Fig. 1a, Table 1). Compared to phyllosphere bacterial strains, soil bacteria were more sensitive to Cd, with EC_5_, EC_50_ and EC_95_ concentrations of approximately 50%, 20% and 10%, respectively, and only about one-fifth of the concentration range (Fig. 1b, Table 1).

**Figure 1.**
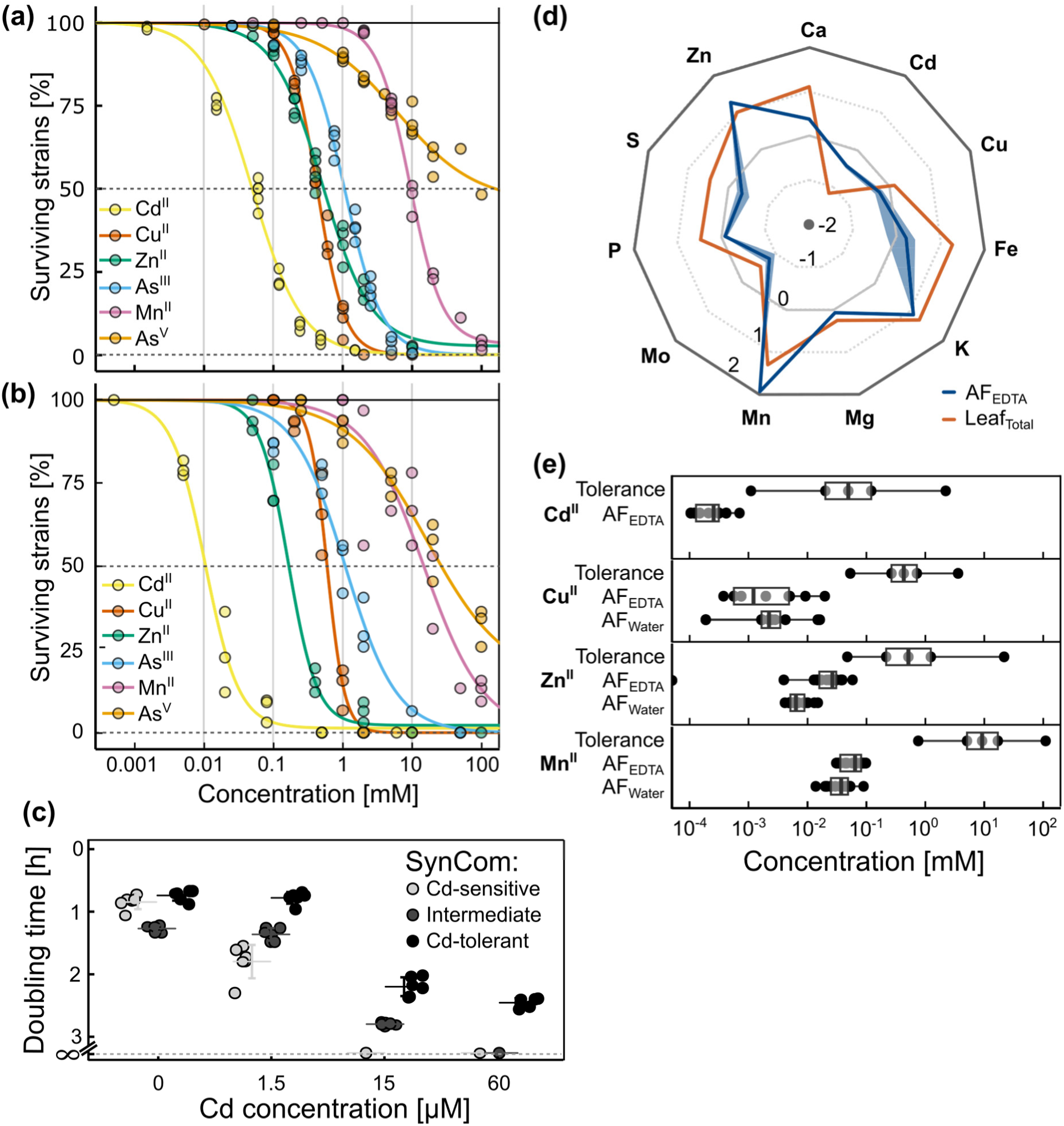
Metal(loid) tolerances of phyllosphere bacterial strains in relation to the leaf apoplastic fluid ionome of *A. thaliana*. **(a, b)** Metal(loid) dose-response curves for the phyllosphere bacterial collection (a, *At*-LSPHERE, *n* = 224 strains) and for a bacterial strain collection from unplanted soils (b, *n* = 33) (Bai et al. 2015). Each strain was spotted on agar-solidified R2A medium supplemented with a series of metal(loid) concentrations. Replicate data points represent independent experiments; lines are curve fits (see Methods, Dataset S1). **(c)** Cd tolerance of synthetic bacterial communities (SynComs; datapoints shown by circles, bars mark mean ± SD, *n* = 6 technical replicates; data from one experiment representative of a total of four independent experiments). Apparent doubling times of the SynComs were calculated based on the earliest identifiable log phase of growth from time series of OD_600_ measurements with cultivation at 26°C for 24 h (see Fig. S3). ∞: no growth. Note that the horizontal axis is not scaled. **(d)** Ionome of leaf apoplastic fluid (AF_EDTA_, blue line), and of the entire fresh leaf (Leaf_Total_, red line). Shown is the average (line) and the standard deviation (shadow) for each element as Log_10_ of the concentration ([mM] for Ca, S, P, Mg, and K; [µM] for Zn, Mo, Mn, Fe, Cu, and Cd; *n* = 4 samples for AF and *n* = 3 samples for leaves). **(e)** Boxplots showing metal concentrations in apoplastic fluid (AF_EDTA_, AF_Water_: extracted in 100 µM EDTA and in water, respectively) of soil-grown plants, alongside metal tolerance of the *At*-LSPHERE collection (re-plotted from (a)). For AF, boxes show median (central line), 25%/75%ile (box), and maximum/minimum (bars) of all measurements (*n* = 17; see Table S5, Fig. S4e-g). For metal tolerances, boxes show effective concentration EC_50_ (central line), EC_25_/EC_75_ (boxes) and EC_1_/EC_99_ (bars)(see Table 1, and Materials and Methods). Dose-response curves are from a minimum of 3 independent experiments conducted on R2A medium (a, b). AF data are from one representative experiment (d; Table S5, GHs 4; Fig. S4e-g, E4) and from 4 independent experiments conducted on greenhouse soil (e). Not shown: AF_Water_ for Cd and data for As were below our detection limits (d, e).

**Table 1.** Effective concentrations of metals for the Arabidopsis phyllosphere (*At*-LSPHERE) and associated soil bacterial strain collections. (based on curve fits).

| *At*-LSPHERE bacterial strain collection |  |  |  |  |
| --- | --- | --- | --- | --- |
| Element | EC<sub>5</sub><br>[mM]<sup>a</sup> | EC<sub>50</sub><br>[mM]<sup>a</sup> | EC<sub>95</sub><br>[mM]<sup>a</sup> | Concentration<br>range<sup>c</sup> |
| Cd<sup>II</sup> | $4.3 \times 10^{-3}$ | 0.049 | 0.56 | 130 |
| Cu<sup>II</sup> | 0.11 | 0.42 | 1.6 | 15 |
| Zn<sup>II</sup> | 0.046 | 0.50 | 5.6 | 120 |
| As<sup>III</sup> | 0.17 | 1.1 | 6.6 | 40 |
| Mn<sup>II</sup> | 1.9 | 9.2 | 46 | 25 |
| As<sup>V</sup> | ~0.32<sup>b</sup> | 147<sup>b</sup> | > 150<sup>b</sup> | >> 100<sup>b</sup> |
| Soil bacterial strain collection |  |  |  |  |
| Element | EC<sub>5</sub><br>[mM]<sup>a</sup> | EC<sub>50</sub><br>[mM]<sup>a</sup> | EC<sub>95</sub><br>[mM]<sup>a</sup> | Concentration<br>range<sup>c</sup> |
| Cd<sup>II</sup> | $2.0 \times 10^{-3}$ | 0.010 | 0.054 | 27 |
| Cu<sup>II</sup> | 0.25 | 0.60 | 1.4 | 5.6 |
| Zn<sup>II</sup> | 0.040 | 0.16 | 0.66 | 17 |
| As<sup>III</sup> | 0.087 | 1.1 | 14 | 160 |
| Mn<sup>II</sup> | 0.77 | 15 | > 100<sup>b</sup> | >> 100<sup>b</sup> |
| As<sup>V</sup> | ~0.40<sup>b</sup> | 27<sup>b</sup> | > 100<sup>b</sup> | >> 100<sup>b</sup> |

<sup>a</sup>EC<sub>*n*</sub> denote metal concentration preventing growth of *n*% of the tested bacterial strains, based on curve fits (see Fig. 1a and b, Dataset S1, and Methods).
<sup>b</sup>Values estimated from experiments (no curve fit possible).
<sup>c</sup>Defined here as EC<sub>95</sub>:EC<sub>5</sub> ratio.

We observed EC_50_ values of 0.42, 0.5, and 1.1 mM, respectively, for the trace elements Cu^II^, Zn^II^, and arsenite. Of these three elements, Zn began to impede the growth of some phyllosphere bacterial strains at the lowest concentration (EC_5_ of 46 µM), and it continued to permit the growth of the most tolerant strains up to the highest concentration (10 mM; Dataset S1, Fig. 1a, Table 1), with a concentration range of 120, very similar to that for Cd. Soil bacteria were less Zn-tolerant, showing EC_5_, EC_50_ and EC_95_ concentrations of approximately 90%, 30% and 12%, respectively, and only about one-seventh of the concentration range, compared to phyllosphere bacteria (Fig. 1b, Table 1).

The 15-fold difference in Cu concentration between the EC_5_ of 0.11 and the EC_95_ of 1.6 mM was the narrowest among all metal(loid)s tested in phyllosphere bacteria. Compared to phyllosphere bacteria, soil bacteria were more Cu-tolerant, with approximately 2-fold EC_5_, 1.5-fold EC_50_, and a similar EC_95_, despite a narrower concentration range of approximately one-third of that observed in phyllosphere bacteria. Arsenite was less toxic to phyllosphere bacteria than Cd^II^, Cu^II^, and Zn^II^, with an EC_50_ of 1.1 mM (Fig. 1a, Table 1).

The EC_50_ of Mn^II^ was as high as 9.2 mM, and arsenate was the least harmful to *At*-LSPHERE strains. At the highest tested arsenate concentration of 100 mM, about 50% of the bacterial strains were still able to grow, although we depleted phosphate in these media (Tables S1 to S3). For As and Mn, phyllosphere bacteria tended to have higher EC_5_ concentrations than soil bacteria (about 2-fold for As^III^ and 4-fold for Mn^II^), similar EC_50_ concentrations, and lower EC_95_ concentrations, clearly different from Zn, Cd and Cu (Fig. 1a and b, Table 1). Taken together, these results suggested that bacterial strains with elevated Cd^II^ and Zn^II^ tolerance are enriched in the phyllosphere of Arabidopsis compared to unplanted soil under natural field conditions.

We next investigated whether the metal tolerances of single strains are representative of those of bacterial communities, using Cd tolerance as an example. We designed three synthetic communities (SynComs) comprising strains that individually exhibited high, intermediate and low Cd tolerance, respectively (Table S4). The composition of these SynComs matched the phylogenetic profile of phyllosphere communities as closely as possible (Fig. S1). On solid R2A media, growth of the SynComs reflected that of the most tolerant strain in single-strain assays (Fig. S2a and S2b). In liquid R2A media, the relative Cd tolerances of SynComs also qualitatively followed those observed for single strains on solid media, but Cd toxicity was more severe in liquid media (Figs. 1c and S3, compare with Fig. S2 and Table 1). These results supported the relevance of our single-strain tolerance assays for exemplary bacterial communities.

### The magnitudes of apoplastic Cd and Zn concentrations reach levels toxic to a subset of phyllosphere bacterial strains

To assess whether phyllosphere bacteria might encounter toxic metal levels in the leaf endosphere, we analyzed the composition of the leaf apoplastic fluid (AF). We cultivated *A. thaliana* in standard greenhouse soil or in a field-collected soil from a region known for geogenically slightly elevated total Cd levels (Stein *et al*., 2017) for 7 to 8 weeks, and extracted apoplastic wash fluid (AWF) using the infiltration-centrifugation technique (Tables S5 and S6) (O’Leary *et al*., 2014; Figueiredo *et al*., 2021). AWF is commonly obtained using water as an infiltration fluid (IF). The IF might affect metal solubility during the experimental procedure, for instance by increasing the extracellular pH within infiltrated leaves. Therefore, we interpret the corresponding results as estimates of the minimal metal levels encountered in the AF. Cell walls contain bound metal cations, which continuously equilibrate with the surrounding AF and can thus be taken up by microbes. To obtain estimates of the maximal metal concentrations in the AF, we used as the IF a solution of 100 µM ethylenediaminetetraacetic acid (EDTA). EDTA is a synthetic chelator of metal cations, with affinities following the sequence of Fe^3+^ > Cu^2+^ >> Zn^2+^ > Fe^2+^ > Mn^2+^ >> Mg^2+^ > Ca^2+^ (Norvell, 1991), expected to mobilize the readily available fraction of transition metal cations from cell walls and other apoplastic binding sites.

Carrying out the infiltration-centrifugation protocol with insufficient caution can damage cell integrity, thus causing cytoplasmic contamination of the AWF. Therefore, we generated additional control samples in which we provoked cytoplasmic contamination by intentionally inflicting mechanical stress on the leaves during the experimental procedure at three levels of severity (little, intermediate, and strong). A principal component analysis (PCA) of the ionomic profiles of all AWF samples demonstrated that provoked cytoplasmic contamination is associated with strongly elevated concentrations of calcium (Ca), potassium (K), magnesium (Mg), sulfur (S), and phosphorus (P) (Fig. S4a). Consequently, a PCA of ionomic profiles is a sensitive means to identify cytoplasmically contaminated AWF samples, all of which were excluded from further analysis (Fig. S4a and B).

Compared to total concentrations across entire hydrated leaves, nutrients (Fe, Mo, Cu, Ca, S, P, Mg, and K) were generally depleted in AF, most strongly for Fe (Fig. 1d, see also Fig. S4c and d). By comparison to bulk leaves, there was an enrichment in AF of Mn, Cd and Zn, consistent with the preferential sequestration of these elements in the apoplastic compartment. Overall, only a minor proportion of the total amount of all elements was extracted from leaves in the AWF (Table S7). We calculated Cd and Zn concentrations in AF_EDTA_ to be at least 0.07 and 5.2 µM, and up to 0.4 and 40 µM, respectively, with considerable variability between replicate samples and independent experiments (Table S5, Fig. 1e, Fig. S4e-g). Accordingly, the range of Zn concentrations in the AF included concentrations capable of preventing the growth of the most sensitive phyllosphere bacterial strains when tested on solid R2A medium. Cd concentrations in AF closely approached these magnitudes (Table 1; Fig. 1E), although plants were grown on greenhouse soil containing only trace levels of Cd (total Cd < 0.1 mg kg^-1^; Table S6). Cd and Zn concentrations in AF were elevated in the presence of EDTA in the IF, suggesting that EDTA either prevented immobilization or mobilized pools of both metals from extracellular binding sites within infiltrated leaves (Fig. 1e, Table S5, Fig. S4a). It should be noted that we consider the direct quantitative comparison between metal concentrations in AF and R2A medium merely as a first reasonable approximation (see also Discussion).

In contrast to Cd and Zn, total concentrations of Cu and Mn in the AF of approximately 1 and 70 µM, respectively, were clearly below toxic levels in the R2A medium between 10 and 100 µM Cu^II^ and 1 to 2 mM Mn^II^ (Dataset S1, Fig. 1a, Table 1). Overall, Cu concentrations in AF were independent of whether we used EDTA solution or water as IF, suggesting that negligible proportions of the apoplastic Cu pool were reversibly bound to cell walls of infiltrated leaves (Fig. 1d, Fig. S4g, Table S5). In summary, these results indicated that Cd and Zn are enriched in the leaf apoplast and have the potential to directly attenuate the growth of a subset of phyllosphere bacteria.

### Towards extrapolating how metals affect phyllosphere bacterial community composition

Previous work demonstrated that the absence of single bacterial strains, such as L203-*Microbacterium*, L231-*Sphingomonas*, L233-*Rhodococcus,* and L262-*Rhizobium*, can alter the structures of bacterial communities (Carlström *et al*., 2019). In these so-called keystone strains, maximum permissive concentration (MPC) was often larger or equal to *At*-LSPHERE-wide EC_50_, however, with strain-specific sensitivities to individual metal(loid)s, so that none of these strains shared an identical profile with any other strain (Fig. 2a, Dataset S1). This suggested that metal(loid) tolerances in keystone strains reflect distinct evolutionary histories rather than shared responses to chemical stress. L233 was the most Cd-sensitive, with an MPC of 1.5 µM Cd (compare Fig. 2a, Table 1), for instance. Although the maximal Cd concentration in AF (0.4 µM) was lower by comparison (see Fig. 1e, Table S5), we observed in liquid R2A medium that a concentration of 0.5 µM Cd suppressed the growth of L203, L231 and L233 (Table S8 and S9).

**Figure 2.**
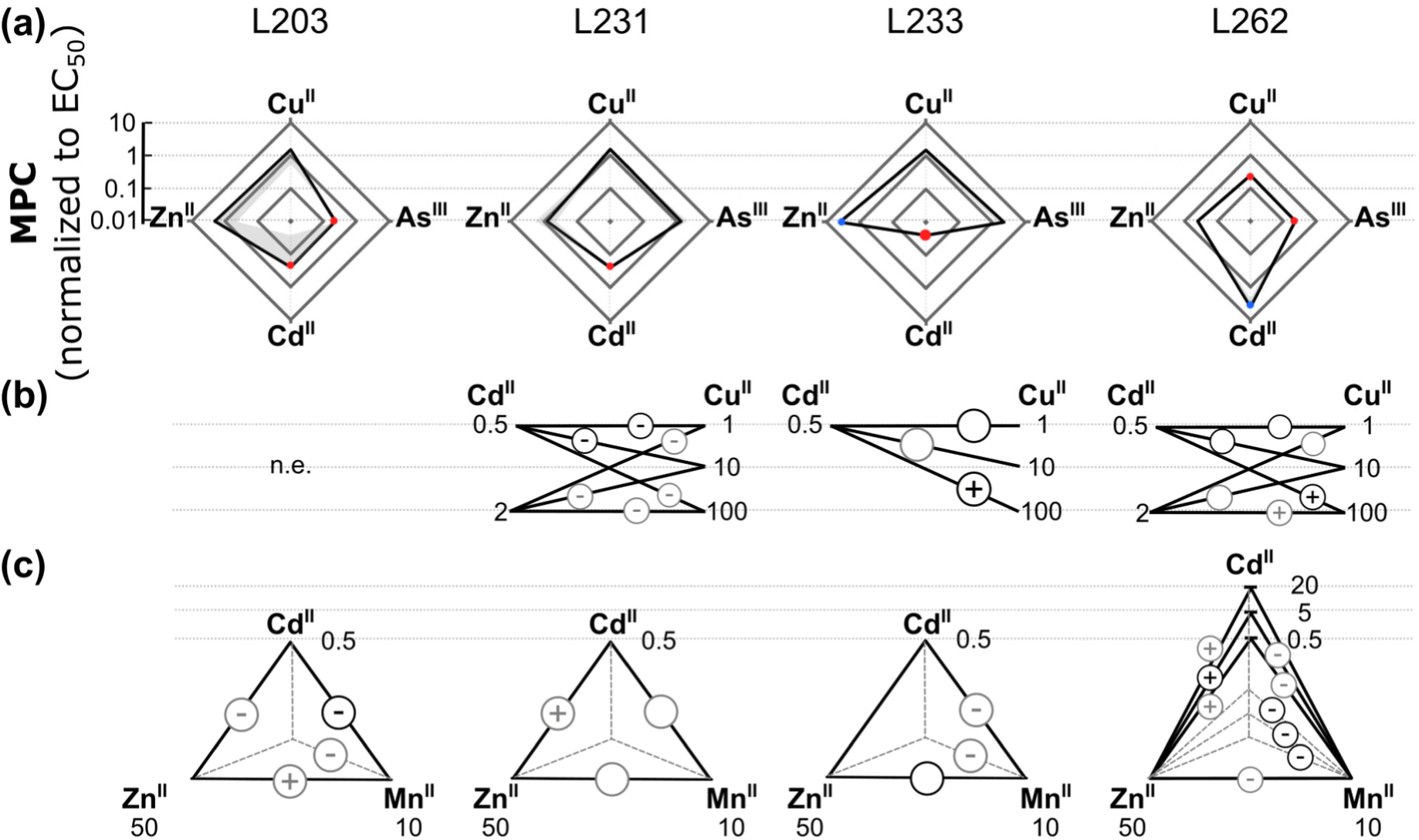
Metal(loid) tolerance profiles of keystone strains of the *A. thaliana* phyllosphere bacterial community. **(a)** Profiles of maximum permissive concentrations (MPC) normalized to EC_50_ on solid R2A media. Shown are median values (connected by black lines), with minima and maxima (grey shadows, *n* = 3 independent experiments), on a Log_10_ scale (1 reflects an MPC equivalent to EC_50_, see Table 1, Dataset S1). Values below 0.5 (small red)/0.05 (large red) and above 5 (small blue) are shown as colored circular shapes. Mn is not shown because of its low toxicity in these strains. **(b, c)** Combined toxicity effects of Cd and Cu (b), and Cd, Zn, and Mn (c) on keystone strains in liquid R2A media (see Tables S8 and S9). Symbols show combined toxicity-enhancing (+) and toxicity-alleviating (-) effects of the metals connected by the corresponding black line; numbers are concentrations [µM]. Results reproduced in three/two out of three independent experiments are shown in black/grey color. Grey dotted lines: effect of the adjacent metal on the combined toxicity of the other two metals (c). Data not shown (n.e.: results could not be evaluated because of metal sensitivity, b). Circle sizes adjusted to fit.

Next, we tested for possible combined effects of metals on the keystone strains. We examined the effects of combining Cd^II^ and Cu^II^, and of combining Cd^II^, Zn^II^, and Mn^II^ in the liquid culture system (Fig. 2b and c). A mild growth reduction appeared at concentrations of 0.5 µM Cd^II^ or 100 µM Cu^II^, whereas the combination of 0.5 µM Cd^II^ and 100 µM Cu^II^ completely eliminated the growth of L233, thus implying synergistic toxicity of Cd^II^ and Cu^II^ (Fig. 2b, Table S8). Similarly, neither Cd^II^ nor Cu^II^ alone were heavily toxic to L262 at the tested concentrations, whereas the combination of 100 µM Cu^II^ with any of the tested Cd^II^ concentrations exacerbated the toxic effects. In contrast, combined treatments appeared to alleviate toxicity in L231 when compared to treatments with Cd^II^ and Cu^II^ alone (Fig. 2b, Table S8). Reproducible combined effects were largely restricted to Cu^II^ concentrations in liquid R2A media that were 14- to 300-fold higher than in AF (see Table S5).

In experiments combining up to three metals, the single Zn^II^ concentration tested (50 µM) alone tended to decrease the growth of L203 and L231, and more strongly of L233 and L262 (Table S9). The tested Mn^II^ concentration (10 µM) alone had no consistent effects on the growth of keystone strains (Table S9). When tested in combination with varying concentrations of Cd^II^ (0.5, 5 and 20 µM), Zn^II^ exacerbated the toxicity of all Cd^II^ conditions in strain L262, which was highly tolerant to Cd alone, whereas Mn^II^ alleviated both the single and combined toxicities of Zn^II^ and Cd^II^ in this strain (Fig. 2c). Zn^II^ toxicity tended to increase in combination with Cd^II^ in L231 and in combination with Mn^II^ in L203. Overall, toxicity-alleviating effects of Mn^II^ tended to predominate, including its effects on L233 and L203 suffering from Cd^II^ toxicity (0.5 µM) or exposed to a combination of Cd^II^ and Zn^II^, but the sizes of these effects were mostly small and insufficient to rescue growth of the same strains at the next higher tested concentration of 5 µM Cd, for instance (Table S9). In summary, these experiments suggested that combined effects are unlikely to cause major increases in metal tolerances of strains compared to single metals as shown in Fig. 1a.

### Phylogeny at the genus level explains most of the variation in metal tolerance profiles

Next, we examined metal tolerance profiles in relation to the phylogeny of *At*-LSPHERE strains. *Actinobacteria* were significantly depleted by Cd concentrations of ≥ 15 µM and enriched among the strains that continued to grow at ≥ 0.2 mM Cu and ≥ 0.4 mM Zn, when compared to other classes (Fig. 3). Relative to other classes, Gammaproteobacteria were enriched at high levels of ≥ 2.5 mM arsenite. Hierarchical clustering of MPCs identified 10 distinct clusters of bacterial strains sharing similar metal(loid) tolerance profiles (Fig. 4).

**Figure 3.**
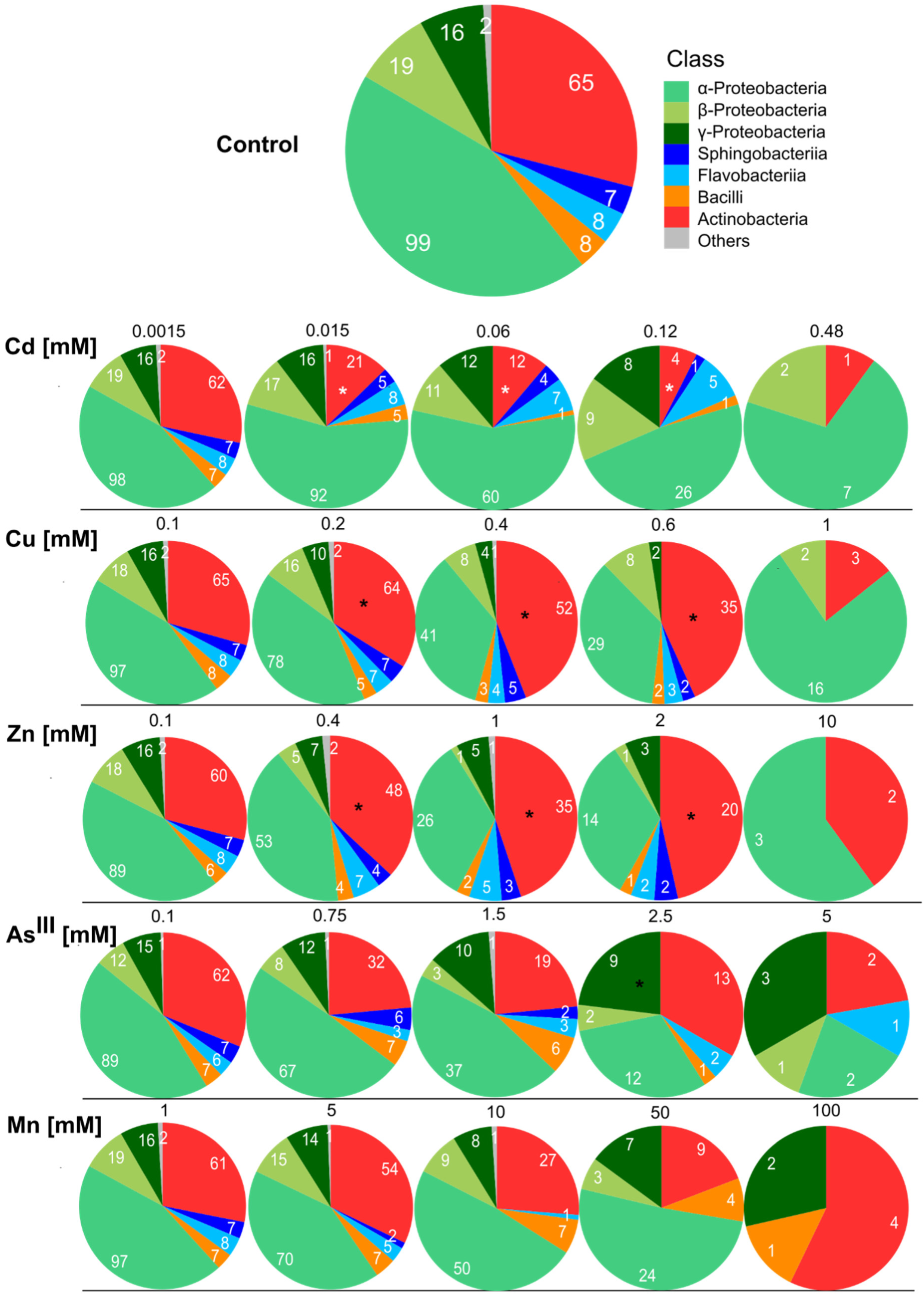
Effects of metal(loid) tolerances of *At*-LSPHERE bacterial strains on the phylogenetic structure among the survivors. Pie fractions reflect the percentage share (with the number given in white) out of all bacterial strains that were able to grow under a given condition (Dataset S1). The metal(loid) concentration is given in black fonts above the chart for each treatment. Asterisks mark significant enrichment (*, black) or depletion (*, white) compared to the control condition (Fisher’s exact test, *q-value* < 0.05).

**Figure 4.**
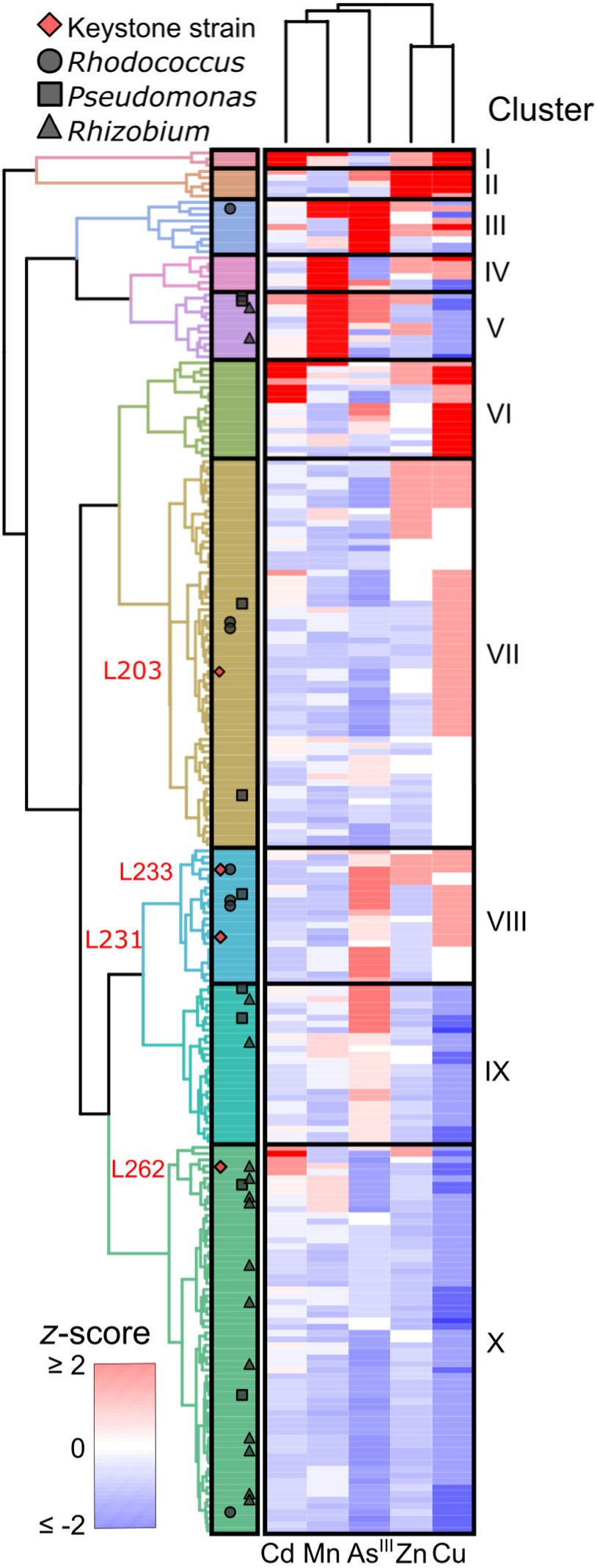
Hierarchical clustering of strains according to their metal(loid) tolerance profiles. Maximum permissive concentrations of Cd, Zn, Cu, arsenite, and Mn in the *At*-LSPHERE strains (Dataset S1) were clustered by Euclidean distance-based (Ward) clustering (*z*-score-based). Clusters (I to X) are emphasized by different colors (left) and separated by horizontal lines. Symbols mark the positions of *Rhodococcus (n* = 6), *Pseudomonas* (*n* = 9), *Rhizobium* (*n* = 15), and keystone strains (*n* = 4).

Cluster I comprised only three strains, L92*-Methylobacterium*, L396*-Bradyrhizobium*, and L401*-Bradyrhizobium* (all α-Proteobacteria), which were highly tolerant to multiple metals but not to the metalloid arsenite. These strains were the most Cd-tolerant (1.5 mM) and tolerated more than twice the EC_50_ concentrations of Cu, Zn, and Mn (Fig. 4, Dataset S1). Cluster II consisted of five highly Zn- and Cu-tolerant strains. Strains highly tolerant to arsenite grouped in cluster III, highly Mn-tolerant strains in cluster IV, and highly Mn-tolerant and Cu-sensitive strains in cluster V. Cluster VI comprised highly to moderately Cu-tolerant strains, some of which were also highly Cd-tolerant. Clusters VII and VIII contained strains with intermediate Cu tolerance, whereby cluster VII contained arsenite-sensitive and cluster VIII contained moderately arsenite-tolerant strains. The final two clusters, IX and X, grouped strains that were highly Cu-sensitive and sensitive to most other metal(loid)s, with intermediate tolerance to arsenite in cluster IX and sensitivity to arsenite in cluster X. Metal tolerance profiles were similar within the genus *Rhizobium*, of which approximately 73% grouped in Cluster X, and 13% in clusters V and IX, respectively. In contrast, the seven *Rhodococcus* strains were distributed into four different clusters: one strain in cluster III, two strains in the neighboring cluster VII, three strains in cluster VIII, with two directly next to one another and a third one further away, and finally one strain in cluster X. Similarly, the nine *Pseudomonas* strains were spread across the five clusters V and VII to X, whereby only two of them clustered next to one another in cluster V. These patterns suggested that phylogenetic relationships account for only part of the variation in metal tolerance profiles.

To obtain more quantitative information on the contribution of phylogeny to the variation in metal tolerance we employed dimensionality reduction-based analyses. There were no major differences between strains of different taxonomic classes according to a PCA of strain-wise MPCs (Fig. S6a), and a constrained principal coordinate analysis (CPCoA) constrained by class explained only 6% of the variance (Fig. S6b*; p <* 0.05). By contrast, CPCoA attributed 42% of the variance to genus (Fig. S6c; constrained by genus, *p* < 0.001). In agreement with this, a representation of metal tolerances on a phylogenetic tree based on 16S rRNA DNA sequences (Bai et al. 2015) suggested that closely related strains mostly exhibited similar metal tolerance profiles (Fig. 5). For example, *Methylobacterium* strains L90, L92, L119, L121, L122, and L123 were highly Cd tolerant, and *Methylobacterium* L90, L92, and L122 were Cu- and Zn-tolerant (Fig. 5). Similarly, all *Bacillus* strains were sensitive to Cd and Cu, and all *Rhizobium* strains (Alphaproteobacteria) were sensitive to Cu and Zn. In summary, metal(loid) tolerance profiles were diverse and partially followed phylogenetic relationships, most prominently at the genus level, suggesting that the remaining phenotypic variation among *At*-LSPHERE strains reflects recent evolutionary divergence.

**Figure 5.**
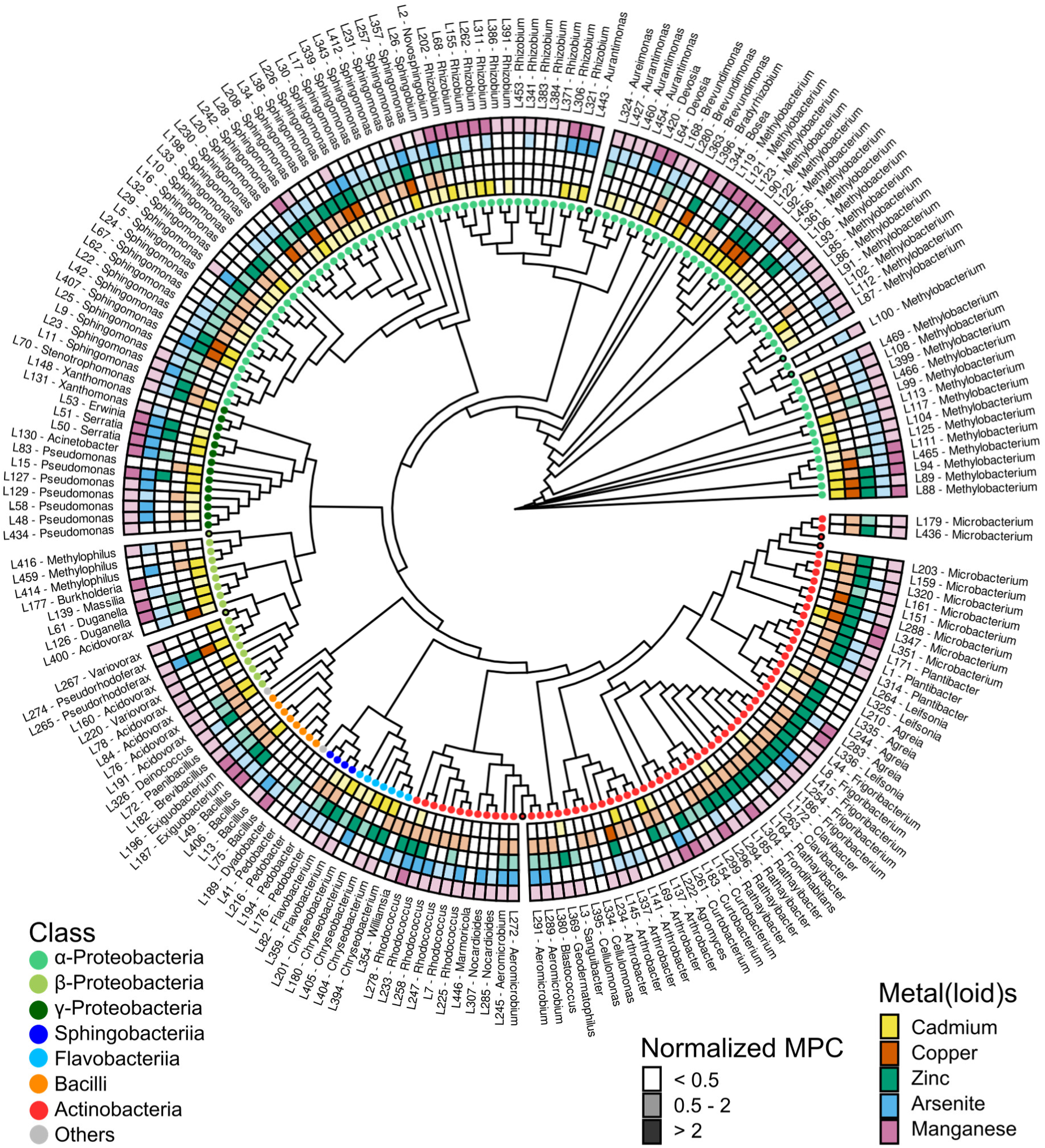
Strain-specific metal(loid) tolerances shown over the phylogeny of the *At*-LSPHERE strain collection. Cladogram of the *At*-LSPHERE strain collection (*n* = 214; 16s rRNA-based (Bai et al. 2015)) with circular heatmap representations of relative tolerance. Heatmap color shade reflects strain-wise maximum permissive concentrations (MPC, median of *n* = 3 independent experiments; Dataset S1) normalized to the EC_50_, the concentration eliminating the growth of 50% of all bacterial strains (Cd: 0.049 mM, Zn: 0.5 mM, Cu: 0.4 mM, As^III^: 1.1 mM, Mn: 9.2 mM; Table 1). Strains (*n* = 8) not profiled for metal tolerance are marked by a black line around the colored circle.

### Genetic basis of differential metal tolerances among bacterial strains

We investigated whether molecular mechanisms that were previously characterized in bacteria can explain between-strain variation in metal tolerances. First, we conducted a PCA of strain-wise MPC for Cd and the orthologs of 9 genes described in the literature to increase Cd tolerance that were present in the genome of each strain (Fig. 6a, Dataset S2). PC 1 clustered the classes *Actinobacteria* and *Bacilli*, which were comparably Cd-sensitive (see Figs. 3 and 5), together towards one side. PC loadings suggested a relationship between enhanced Cd tolerance and the presence of *czcA, czcB, czcC, czcD*, *smtA*, and *tolC*, of *zntA* for PC1 only, as well as the absence of *zntR* and *cadC..smtB* (Fig. 6a). Beyond the identity of Cd tolerance genes, their gene dosage (or copy number) could be expected to scale with strain-level Cd tolerance. A Kendall’s rank correlation analysis of the gene score, which we calculated as the cumulative number of Cd tolerance-related genes and gene copies present in the genome, and Cd MPC showed a significant correlation (*τ* = 0.31, *p* < 3.5 × 10^-9^; Fig. 6b).

**Figure 6.**
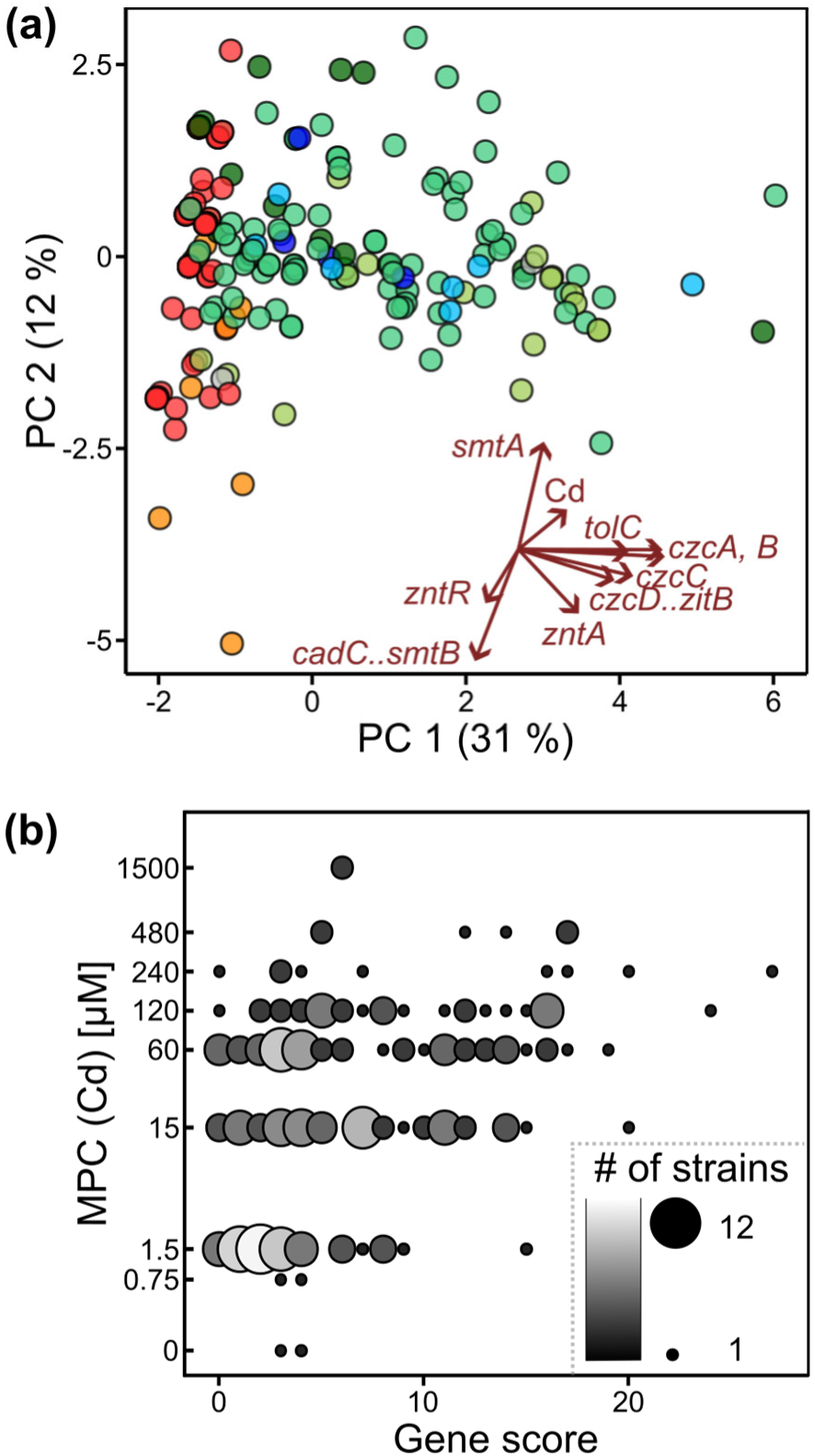
Relationships between known Cd tolerance genes and Cd tolerance of *At*-LSPHERE strains. **(a)** PCA of strain-wise maximum permissive Cd concentrations (MPC, *z*-score-based) and presence/absence of known Cd tolerance-related genes. Each datapoint represents one strain (Datasets S1-S3). The percentages of variation accounted for by each significant PC are given in parentheses (95% confidence intervals: PC 1, 31% to 36.7%; PC 2, 10.9% to 13.8%; bootstrapping with 999 replicates). KEGG Orthology (KO) identifiers with significant loadings on PC1 are: *czcA–D* and *tolC* (alternative short names linked by “..”; Dataset S6). **(b)** Plot of strain-wise maximum permissive Cd concentration (MPC, absolute) against a gene score incorporating both presence/absence and copy number of known Cd tolerance-related genes. Each datapoint represents one strain (Datasets S1-S3). The gene score of a strain is the genome-wide sum of genes and gene copies with previously published roles in Cd tolerance (Kendall’s rank correlation, *τ* = 0.31, *p* = 3.5×10^-9^; Dataset S7).

We next broadened our analysis to include known metal(loid)-related (100 genes of interest, GoI, Dataset S2). CPCoA indicated that 22% of the presence/absence genetic variation for these genes was explained by class (constrained by class, *p* ≤ 0.001), and 63% by genus (constrained by genus, *p* ≤ 0.001) (Fig. S7a-c). This result further highlighted the importance of genus-level taxonomy, consistent with the pattern observed for phenotypic variation in metal tolerances (Fig. S6). By comparing median MPC of all strains containing the GoI in their genomes with that of all strains lacking the GoI (Datasets S3 and S4), we identified 51 genes that were significantly associated with tolerance to one or several metal(loid)s: 21 genes (41%) for Cd, 9 for Cu (18%), 15 (29%) for Zn, none for As^III^, 20 (39%) for Mn, and 9 (18%) for As^V^ tolerance (Table S10, Dataset S4, Wilcox rank-sum test, *p*-values adjusted with Holm-Bonferroni corrections). We detected positive effects for *czcA, czcB, czcC, tolC* and *zntA* on Cd tolerance, in agreement with Fig. 6a, whereby statistical significances were the highest for *czcA*, *czcB* and *tolC* (Dataset S4, Fig. 7, *p_adj_* < 0.005). In addition to genes previously implicated in bacterial Cd tolerance, our analysis implicated *arsC1..arsC*, *arsH*, *cusB*..*silB*, *mdtA*, *mdtB*, *mdtC* and *zur* as candidate genes for increasing Cd tolerance. Conversely, the presence of *comB*, *cmtA*, and *smtB* each decreased the median MPC of Cd by a factor of 40. *Actinobacteria* and *Bacilli* of the *At*-LSPHERE collection were characterized by an overall lack of Cd tolerance genes and the presence of Cd-sensitizing genes, consistent with their Cd sensitivity (Fig. 7, see Figs. 3 to 5 and 6a). Similarly, there was an agreement between Cd tolerance and the genomic content of Cd-related genes identified here in the most Cd-sensitive strains (L454, L172, L263, and L406) and one of the two most highly Cd-tolerant strains (L396) (Fig. 7). As an exception to this pattern, the genome of the highly Cd-tolerant strain L92*-Methylobacterium,* contains only three known Cd tolerance-increasing genes, *czcA, czcB* and *zntA*. Taken together, these results implicate orthologues of previously characterized metal(loid)-related genes in specific metal(loid) tolerances of phyllosphere bacterial strains and suggest further contributions from unknown or poorly annotated genes.

**Figure 7.**
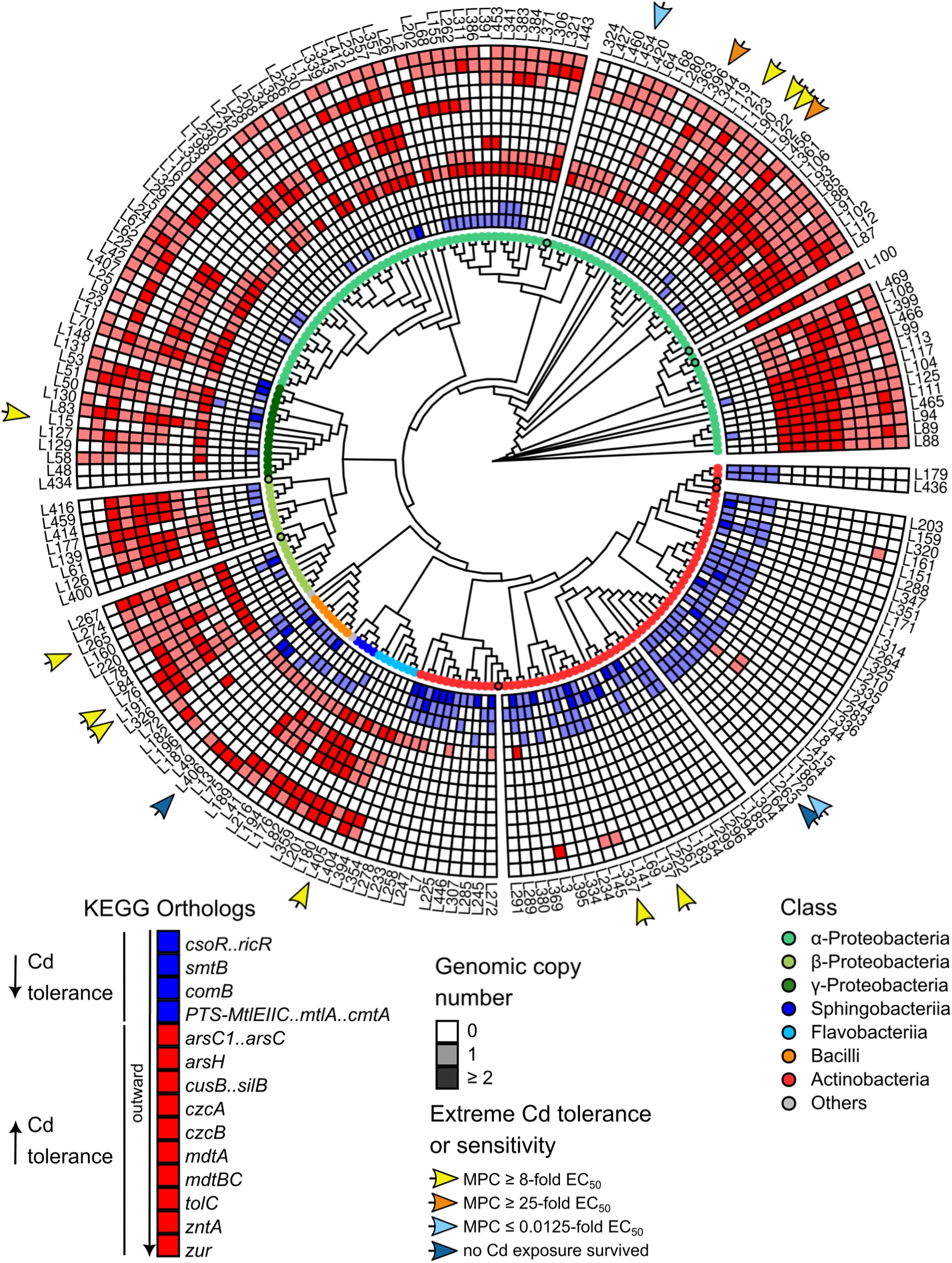
Metal(loid)-related genes implicated in Cd tolerance of *At*-LSPHERE strains shown over phylogeny. Shown are known metal(loid)-related genes of which the presence in the genome was statistically associated with altered Cd tolerance in our study (*p*_adj_ < 0.005, for *arsH p*_adj_ = 0.023; with Bonferroni corrections). Colors of KO identifiers reflect whether the median of maximum permissive concentrations (MPC median) of all positive strains was > 2-fold higher (red) or > 2-fold lower (blue) than that of all strains lacking an orthologue (Dataset S4; else KO id. not shown). Genes (KO identifiers) are only shown if present in at least 30 bacterial strains (Dataset S4). Strains (*n* = 8) not profiled for metal tolerance are marked by a black line around the colored circle.

## Discussion

### Metal tolerances of individual strains and their relevance for bacterial community composition in the leaf endosphere

Our results support the conclusion that the levels of Zn and Cd, and possibly also of other metal(loid)s, in the leaf apoplast contribute to shaping bacterial community composition in the leaf endosphere of *A. thaliana* by eliminating or attenuating the growth of sensitive bacterial genotypes. First, we observed marked overall differences in the degrees of tolerance to Zn and Cd between the bacterial strain collection from the phyllosphere and the collection from unplanted soil (Fig. 1, Table 1). This suggests an enrichment of strains exhibiting elevated tolerance to these metals among in the *At*-LSPHERE collection by comparison to the collection from unplanted soil. Second, Zn and Cd concentrations in the leaf apoplastic fluid (AF) of soil-grown Arabidopsis plants closely approached, or overlapped with, the metal concentrations that were toxic to metal-sensitive bacterial strains of the *At*-LSPHERE collection in solid R2A media (Figs. 1 and S4, Table S5). In addition, Cd, Mn and Zn were enriched in AF compared to their concentrations in bulk leaves. Third, two keystone strains (Carlström *et al*., 2019), L233-*Rhodococcus* and L203-*Microbacterium*, were among the most sensitive strains (Fig. 2 and Fig. S5), implying the potential for pronounced secondary effects of Zn and Cd toxicity on overall species composition in the leaf endosphere. Finally, a set of exemplary tests conducted by us suggested that between-strain and between-metal interactions had no, or only few, and mostly small, reproducible effects alleviating Zn or Cd toxicity (Figs. 1 and 2, S2 and S3, Tables S8 and S9).

Our study has some limitations that should be considered when interpreting these conclusions. While partly shared soil environments of origin support the comparability of the bacterial collections (Bai *et al*., 2015), the number of strains was considerably smaller in the soil collection than in the *At*-LSPHERE collection, thus potentially decreasing the precision of the values summarizing metal(loid) tolerances across soil bacterial strains (Fig. 1). Relating metal concentrations in leaf AF to those in solid or liquid bacterial growth media fundamentally remains an extrapolation. Considerable proportions of the total metal concentrations accessible to and measured in AF, as well as those measured in R2A media, are present in chelated form in solution, within precipitates or bound to the surfaces of solids. Yet, the degree of Cd and Zn toxicity to bacteria is likely to depend on the concentrations of their free aqueous cations. In addition to this, the levels of other solutes can strongly influence metal toxicity. Importantly, other divalent cations compete with toxic metal cations for bacterial uptake, and the levels of bioavailable nutrients, such as phosphate, sulfur, iron and manganese, can alleviate metal toxicity inside bacterial cells. The overall elemental composition of solid R2A media (*ca*. 5-fold Fe, 10% Ca, 30% Mg, 20% K, 6-fold S, and 2.6-fold P, compared to AF) suggested that bacterial metal toxicities are broadly comparable between the two matrices (Table S5). The agar polysaccharide in solid R2A medium contributes a macromolecular matrix of metal cation binding sites, similar to the cell walls of Arabidopsis which interact with the AF *in vivo*. Agar also contributes substantial quantities of contaminant Ca^2+^ and Mg^2+^ cations capable of competing with Cd^2+^ and Zn^2+^, compared to liquid R2A media (Table S3). Thus, unsurprisingly, bacteria generally tolerated higher Cd concentrations in solid than in liquid R2A medium (≥ 16-fold difference for L262 and the intermediate SynCom, > 10-fold for L203; compare Fig. 1c with Fig. S2, Fig. 2a with Suppl. Fig. S5).

We expect the composition of leaf AF to be highly dynamic, which must be kept in mind when comparing with the metal concentrations in our bacterial tolerance assays. In addition to the concentrations of free metal cations and other contributing factors outlined above, a pH decrease (Geilfus, 2017; Geilfus *et al*., 2020) and a more oxidized redox state can directly enhance metal toxicity in AF by decreasing the proportion of metal cations in complexed or bound from, thereby increasing the bioavailable fraction of free aqueous metal cations. Importantly, in response to infection by the necrotrophic fungus *Plectosphaerella cucumerina*, Arabidopsis accumulated Zn^2+^ in the leaf apoplast dependent on the Zn^2+^/Cd^2+^ efflux pump-encoding genes *AtHMA2* and *AtHMA4* (Escudero *et al*., 2022). Thus, microbe-induced fluxes of metal cations across the plasma membrane of plant cells may be more widespread than is currently recognized. Interpreting the differences in overall metal tolerances of the *At*-LSHERE collection compared to soil bacterial communities as an outcome of selective strain enrichment would imply that in phyllosphere bacterial communities even the mildest Zn and Cd toxicities are ecologically relevant (compare EC_5_ in Table 1). We observed the largest relative differences between *At*-LSPHERE and soil bacterial Zn and Cd tolerances for EC_95_ (Table 1). One possible interpretation of these observations could be that the corresponding metal levels are ecologically relevant for Zn^II^ (corresponding to the degree of toxicity inflicted by 0.66 to 5.6 mM Zn in solid R2A medium), Cd^II^ (0.054 to 0.56 mM Cd) and possibly also Cu^II^ (1.4 to 1.6 mM Cu^II^).

In contrast to Zn, Cd has no known nutritional role in Arabidopsis, raising the question of acquisition and partitioning of Cd by plants in conjunction with a possible biological role in plant-microbe interactions suggested by our results. In addition to HMA2 and HMA4, and among other characterized membrane transporters (Sterckeman & Thomine, 2020), Cd^2+^ is a secondary substrate of the high-affinity root Fe^2+^ uptake systems IRT1 of Arabidopsis and RIT1 of pea (Connolly *et al*., 2002; Vert *et al*., 2002; Cohen *et al*., 2004). In the Zn- and Cd-hyperaccumulating species *Arabidopsis halleri* and *Noccaea caerulescens*, extremely high concentrations above 3 mg g^-1^ Zn and 0.1 mg g^-1^ Cd in bulk leaf dry biomass are thought to function as elemental defenses against biotic stress, including bacterial pathogens (Fones *et al*., 2010; Hörger *et al*., 2013; Kazemi-Dinan *et al*., 2014; Stein *et al*., 2017). Maximal Zn and Cd concentrations in leaves of healthy *A. thaliana* plants are less than 10% by comparison, but this does not preclude the possibility of a highly localized accumulation of Zn or Cd to levels that can be toxic for bacteria. Much higher bacterial toxicity of Cd compared to Zn (Table 1), as well as the absence of a correlation between Cd and Zn tolerance among *At*-LSPHERE strains (Fig. 4), argue against the possibility that our findings could be a consequence of the similarity between the chemical properties of Cd and Zn.

For metal(loid)s other than Zn and Cd, there was substantially less evidence for a role in shaping phyllosphere microbial communities. Cu^II^ concentrations in AF were far below Cu^II^ levels toxic for *At*-LSPHERE bacteria on agar-solidified media, and *At*-LSPHERE bacteria were not generally more Cu^II^-tolerant than soil bacteria (Fig. 1). Cd toxicity to the keystone strains L233-*Rhodococcus* and L262-*Rhizobium* was potentiated by the additional presence of Cu^II^ only at a high concentration of 100 µM in liquid R2A medium (Fig. 2). However, some existing knowledge supports that Cu could be relevant in plant-microbe interactions. Human macrophages mediate the local accumulation of Cu ions in the phagolysosome to kill pathogens (White *et al*., 2009; Achard *et al*., 2012; Samanovic *et al*., 2012). Transcript profiling revealed that several copper-related genes were upregulated in *A. thaliana* leaves in response to the strain *Methylobacterium extorquens* PA1 (Vogel *et al*., 2016), and the treatment of Arabidopsis with Cu was reported to enhance its immunity against *Pseudomonas syringae* pv. *tomato* DC3000, for example (Xia *et al*., 2026). The strongest biotoxic effects of Cu generally result from redox cycling between oxidized Cu^II^ and reduced Cu^I^, and from exposure to Cu^I^ (Samanovic *et al*., 2012). Similar to their proposed functions in human immune cells, plasma membrane-localized gp91^phox^ NADPH oxidases, such as the superoxide-producing *At*rbohD and *At*rbohF (Nathan & Shiloh, 2000; Torres *et al*., 2002) and the Cu^II^ chelate reductases *At*FRO4 and *At*FRO5 (Bernal *et al*., 2012), could contribute to redox cycling of Cu in the plant phyllosphere. Indeed, *AtrbohD* is important for the maintenance of microbiota in leaves of *A. thaliana* (Pfeilmeier *et al*., 2021). Total As was below our detection limits in AWF, and bacteria seemed to be comparably tolerant to arsenite (As^III^) and arsenate (As^V^) on solid R2A medium. Organo-arsenicals, which can be far more toxic than inorganic forms of As, warrant further investigation, given that *A. thaliana* can take up dimethylarsenate and convert it into dimethylmonothioarsenate- and dimethyldithioarsenate, for example (Colina Blanco *et al*., 2023). For Mn and As^III^, only EC_5_ values were higher in the *At*-LSPHERE than in the soil bacterial collection, implying a possible ecological relevance associated with mildly toxic concentrations in AF (Table 1), which will require further work. The finding that Mn^II^ alleviated Cd^II^ toxicity in keystone strain L203-*Microbacterium* in liquid R2A media may be related to this (Fig. 2).

To date, the information available from previous metal(loid) tolerance screens of bacterial strain collections is limited. Much of the available data are from single bacteria hypothesized to exhibit enhanced metal(loid) tolerance (Tada & Inoue, 2000; Zhao *et al*., 2015), and from multiple strains collected in environments characterized by high levels of abiotic stress or heavy metals (Nieto *et al*., 1987; Idris *et al*., 2004; Abou-Shanab *et al*., 2007; Barzanti *et al*., 2007; Fones *et al*., 2010; Luo *et al*., 2011; Fones *et al*., 2016). Overall, these studies concur with our results regarding the comparably high toxicity of Cd and the overall narrow concentration range for Cu^II^ (see Table 1, Fig. 1). Although *Actinobacteria* were the least Cd-tolerant among *At*-LSPHERE strains, a study of heavy metal-tolerant *Actinobacteria* identified strains able to tolerate Cd and Cu concentrations of up to 100 mM, i.e., about 70- and 100-fold higher concentrations than the most tolerant strains of the *At*-LSPHERE collection (Amoroso *et al*., 1998). Across these previous studies, natural variation in bacterial metal tolerances is very large, and the environment of bacterial origin appears to be more influential than their phylogeny. Metal tolerances of *At*-LSPHERE strains were mostly lower than those reported in bacteria originating from metal-rich environments, with some remaining uncertainty given the diversity of media and designs among tolerance assays used.

### Relationship of metal(loid) tolerance with phylogeny and the gene content of bacterial genomes

Phylogenetically closely related bacteria generally tended to exhibit similar metal tolerance profiles, and our results supported a genetic basis of differential metal tolerance profiles, despite a few notable exceptions (Figs. 4 to 7), consistent with previous findings among bacterial leaf endophytes of metal hyperaccumulator plants (Fones *et al*., 2010; Fones *et al*., 2016). We identified some candidate genes in *At*-LSPHERE strains among the genes annotated as orthologues of metal(loid)-related based on previous work in bacterial models (Figs. 7, Table S10). Of the genes strongly implicated in Cd tolerance in the *At*-LSPHERE collection, the *czc* and *znt* operons are well-known (Figs. 6 and 7, Table S10, Dataset S4). The CzcCBA efflux system of *Ralstonia metallidurans* pumps Cd out of cells (Nies *et al*., 1989; Nies & Silver, 1989; Nies, 1992b) for Cd resistance (Nies, 1992a; Silver & Walderhaug, 1992), supported by a virtual model of *R. metallidurans* (Legatzki *et al*., 2003). Bacterial ZntA Zn/Cd/Pb-transporting P-type ATPase proteins often function in Zn detoxification and can be relevant for Cd resistance (Beard *et al*., 1997; Rensing *et al*., 1997; Outten *et al*., 1999; Singh *et al*., 1999; Choudhury & Srivastava, 2001; Chaoprasid *et al*., 2015). A low concentration of Cd (50 µM) completely inhibited growth of an *Agrobacterium tumefaciens* C58 loss-of-function *zntA* mutant (Chaoprasid *et al*., 2015). We did not identify the gene encoding the known regulator of *zntA*, *zntR* (Table S10), and its presence even appeared to be inversely related to Cd tolerance (Fig. 6), which might reflect mutual interference between the toxin Cd^2+^ and the micronutrient Zn^2+^.

In addition to known Cd tolerance mechanisms, our data implicated Resistance-Nodulation-Division (RND) type multidrug efflux pump component-encoding *mdtA*, *B*, and *C* in Cd tolerance of *At*-LSPHERE bacteria. In *Escherichia coli*, periplasmic MdtA connects the inner membrane efflux pump MdtBC to the outer membrane channel TolC to confer antibiotic resistance (Nagakubo *et al*., 2002; Nishino *et al*., 2007; Kim *et al*., 2010). The fourth gene of the *mdtABCD* operon *mdtD* encodes a putative transporter MdtD, which is also involved in antibiotic resistance and can promote the efflux of citrate, later renamed iron-citrate efflux Transporter IceT in *Salmonella typhimurium* (Frawley *et al*., 2013). A knockout of *tolC* in *E. coli* resulted in Cd hypersensitivity (Achard-Joris *et al*., 2005). Moreover, mRNA of genes encoding the AcrAB-TolC efflux pump (*acrB* and *tolC*) were upregulated in Cd-adapted *Salmonella* Typhi Ty2 cells (Kaur *et al*., 2021). The MdtABC system was implicated in Zn and Cu tolerance in *Salmonella enterica* (Nishino *et al*., 2007). Supporting the *in planta* relevance of RND-type efflux systems, an RND family efflux transporter MFP subunit (Mext_2832) was detected in the *in planta* proteome of *M. extorquens* PA1 (Müller *et al*., 2016a), and efflux systems including CzcABC family RND transporters were among the proteins induced during phyllosphere colonization in *Rhizobium* L68 and *Sphingomonas* L257 (Hemmerle *et al*., 2022).

Our results implicated genes with known roles in As tolerance, *arsC1..arsC* and *arsH*, in Cd tolerance of *At*-LSPHERE strains (Figs. 6 and 7, Table S10, Dataset S4). Among three different KO identifiers linked to *arsC* (*arsC1*..*arsC*, K00537; *arsC2*..*arsC*; K03741, *arsC*, K18701; Dataset S4), only *arsC1..arsC* (*arsC1*), described as arsenate reductase that uses glutathione as a cofactor (glutaredoxin) in the KEGG database, showed a positive effect on Cd tolerance. Dhankher *et al*. (2003) reported that bacterial *arsC1* conferred Cd tolerance to *E. coli*, and to *A. thaliana* and *Nicotiana tabacum* plants, but the underlying mechanism remains unidentified. The second Cd tolerance candidate gene in the *ars* operon, *arsH* was described as a methylarsenite oxidase that can form the less toxic methylarsenate (Yang & Rosen, 2016) and as an H_2_O_2_-producing enzyme that can alleviate oxidative stress in *Synechocystis* PCC 6803 (Lopez-Maury *et al*., 2003). In *Pseudomonas chenduensis* strain MBR, *arsH* transcript levels were upregulated under Cd toxicity (Li *et al*., 2020). There is no published information on roles in Cd tolerance for the two remaining Cd tolerance candidate genes, *cusB* (annotated to encode a component of a Cu^+^ detoxification system), and *zur* (encoding a Zn sensor mediating the transcriptional repression of genes encoding high-affinity Zn uptake systems in its Zn^2+^-bound form). Effective repression of Zn^2+^ uptake may help to decrease inadvertent uptake of Cd^2+^. We detected no effect for *cadC..smtB*, which was previously implicated in bacterial Cd tolerance (Endo & Silver, 1995).

Future work should test further the hypothesis supported by our results that apoplastic Zn and Cd contribute to the structuring of bacterial communities in the leaf endosphere of *A. thaliana*, as well as investigate the roles of physiologically relevant chemical forms of Cu and As that could not be tested here. What currently remains limiting is the lack of cell biology tools for quantitatively monitoring the dynamics in apoplastic free metal cation concentrations, and of genetic tools for changing these without affecting other aspects of plant metal homeostasis that could have indirect effects on microbial performance. The future exclusion of epiphytic bacteria may facilitate the analysis of interactions between the plant host and bacteria in the leaf endosphere (Chen *et al*., 2020). Finally, functional studies can now target metal(loid) tolerance candidate genes identified here in the published genomes of *At*-LSPHERE bacteria.

## Acknowledgements

We thank Prof. Dr. Julia Bandow for access to 96-pin microplate replicator. We thank Andreas Aufermann, Rebekka Fresen and Martin Pullack for assistance with plant cultivation and Petra Düchting for contributing her expertise and for conducting all multi-element analyses.

## Competing Interests

The authors declare no competing interests.

## Author Contributions

U.K. conceived the project, J.F.P.-M. and U.K. designed research with contributions from J.A.V., J.F.P.-M. and A.T. performed experiments, J.A.V. contributed materials, J.F.P.-M., A.T., and U.K. analyzed data, J.F.P.-M. and U.K. wrote the manuscript, with contributions from J.A.V. All authors read and edited the manuscript.

## Data Availability

The data associated with this manuscript are provided in the manuscript or in the Supporting Information. Any additional raw data will be made available by the authors upon reasonable request.

## Supporting Information

**Dataset S1.** Maximum permissive metal concentrations.

**Dataset S2.** KEGG orthologous gene groups and genes of interest (GoI).

**Dataset S3.** Genomic presence/absence of genes based on Kegg Orthologue (KO) identifiers.

**Dataset S4.** Effects of previously described metal-related genes on tolerance levels (median MPC) in At-LSPHERE strains.

**Dataset S5.** Elemental concentrations and glucose-6-phosphate dehydrogenase (G6PDH) activity in AWF in a set-up experiment.

**Dataset S6.** Loadings of genes on principal components (Figure 6A).

**Dataset S7.** Genomic presence/absence and copy numbers of Cd-related genes and gene scores (Figure 6B).

**Figure S1.** Taxonomic structure of synthetic communities.

**Figure S2.** Cd tolerance of synthetic bacterial communities and single strains on agar-solidified R2A media.

**Figure S3.** Cd tolerance of synthetic bacterial communities based on growth curves in liquid R2A media.

**Figure S4.** Ionomes of leaf and leaf apoplast samples of *A. thaliana*.

**Figure S5.** Cd tolerance of keystone strains in liquid media.

**Figure S6.** Relationship between metal tolerance profiles and phylogeny of *At*-LSPHERE strains.

**Figure S7.** Relationship between metal-related genes in the genome and phylogeny of *At*-LSPHERE strains.

**Table S1.** Composition of R2A medium.

**Table S2.** Metal salts used in tolerance screening on solid R2A medium.

**Table S3.** Total concentrations of elements in R2A media as quantified by ICP-OES.

**Table S4.** Cd tolerances of individual strains of the synthetic communities.

**Table S5.** The ionome of the *A. thaliana* leaf apoplastic fluid.

**Table S6.** Soil ionomes for the greenhouse soil and the field-collected soil mixtures used for plant cultivation.

**Table S7.** Leaf ionomes before and after AWF extraction for *A. thaliana* grown in greenhouse soil or field soil mixture.

**Table S8.** Combined effects of cadmium and copper toxicity on keystone strains.

**Table S9.** Combined effects of cadmium, zinc and manganese on keystone strains.

**Table S10.** Known metal-related genes associated with metal tolerances in the *At*-LSPHERE collection.

## Notes

### Competing Interest Statement

The authors have declared no competing interest.

### Summary of Updates

shortened, small errors corrected in various places, formatting altered

### Funding

- Deutsche Forschungsgemeinschaft: RTG 2341 Microbial Substrate Conversion (MiCon)
- Ruhr University Bochum: no number

