## Supplemental Figures, Tables and Methods text for "Metal toxicity contributes to the structuring of bacterial communities in the Arabidopsis phyllosphere"

This PDF file includes:

Figure S1 to S7

Tables S1 to S10

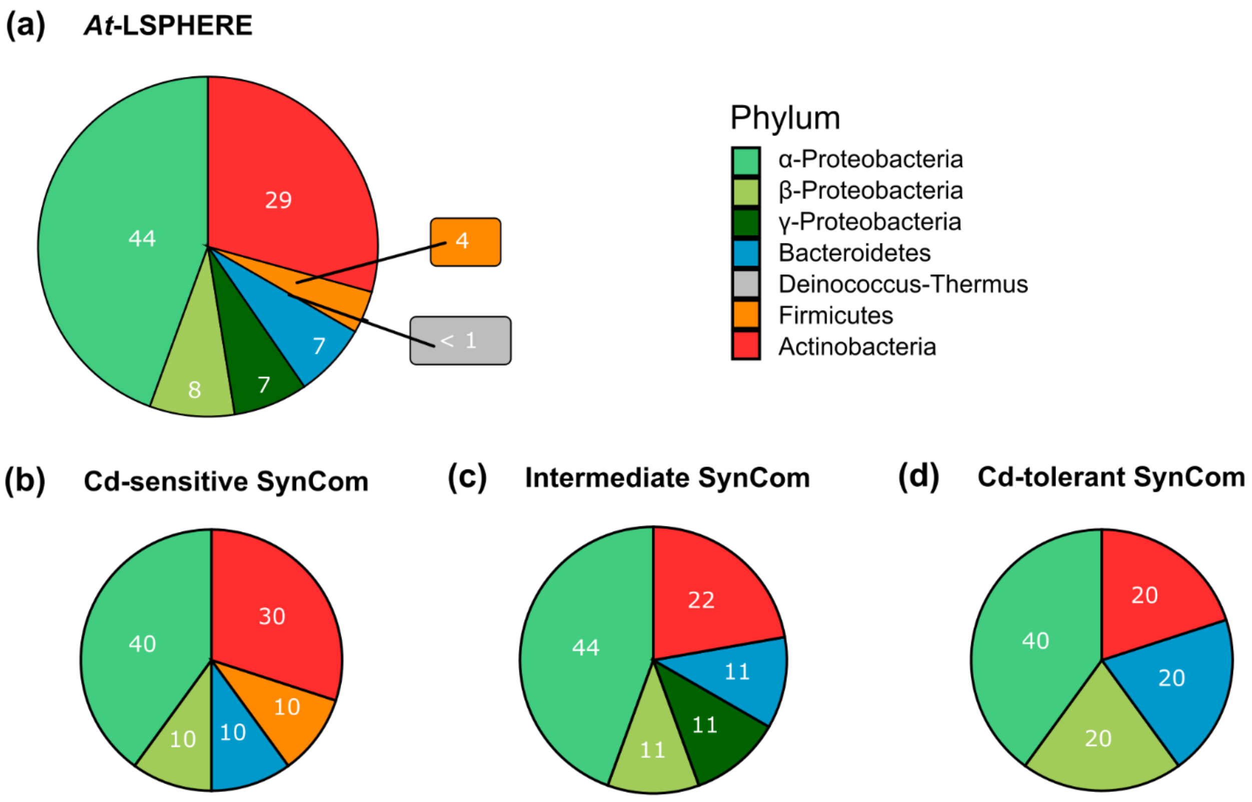


**Figure S1. Taxonomic structure of synthetic communities. (a-d)** Taxonomic structures of the full *At*-LSPHERE collection ((a), *n* = 224), the Cd-sensitive synthetic community ((b), *n* = 10), the intermediate SynCom ((c), *n* = 9), and the Cd-tolerant SynCom ((d), *n* = 10). Bacterial classes are color-coded; strains of each SynCom are specified in Figure S2(b) and Table S4.

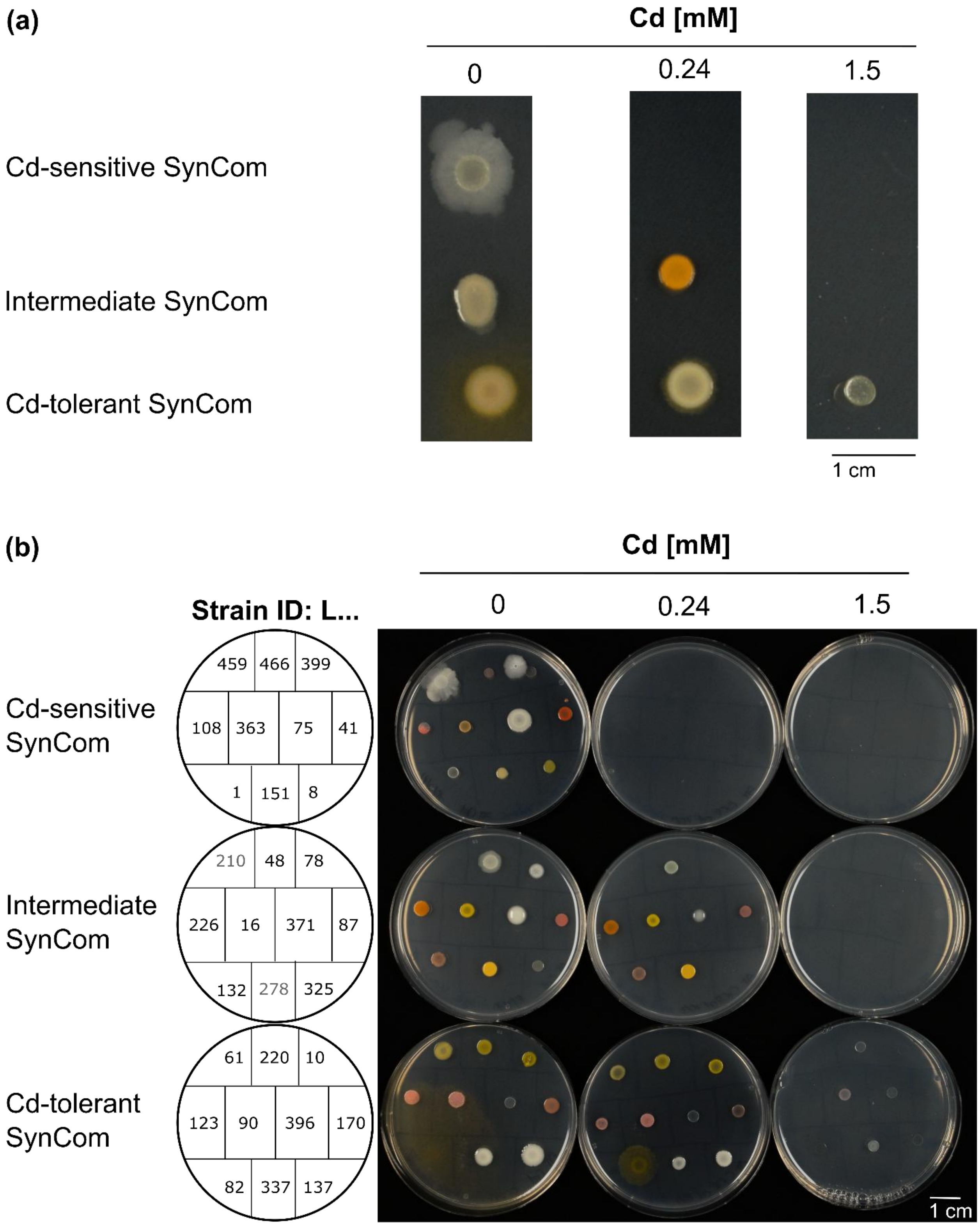


**Figure S2. Cd tolerance of synthetic bacterial communities and single strains on agar-solidified R2A media. (a, b)** Photographs of the Cd-sensitive, intermediate, and Cd-tolerant SynComs (a) and of all single strains of each SynCom (b) spotted on agar-solidified R2A media supplemented with Cd (0, 0.24 and 1.5 mM Cd), after cultivation at RT for 10 d.

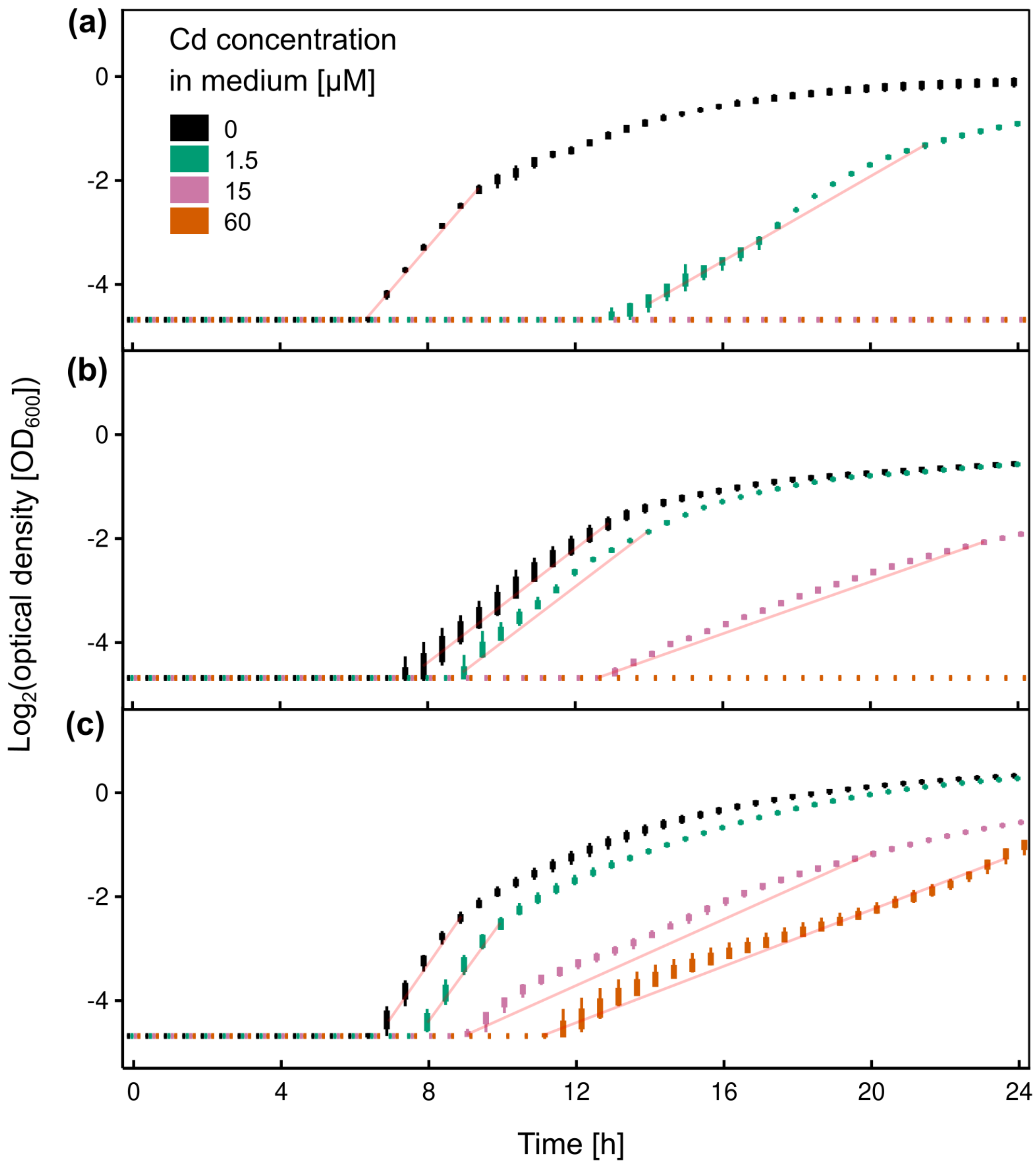


**Figure S3. Cd tolerance of synthetic bacterial communities based on growth curves in liquid R2A media. (a-c)** Shown are growth curves as time series of optical densities of SynComs cultivated in liquid R2A media containing 0, 1.5, 15, and 60 µM Cd for the Cd-sensitive (a), intermediate (b) and Cd-tolerant (c) SynCom (boxplots of $n = 6$ technical replicates). Red lines mark the range of datapoints that were used to calculate doubling times shown in Figure 1c. Data are from one experiment representative of a total of four independent experiments. The optical density was measured at 600 nm once per h during cultivation at 26°C for 24 h.

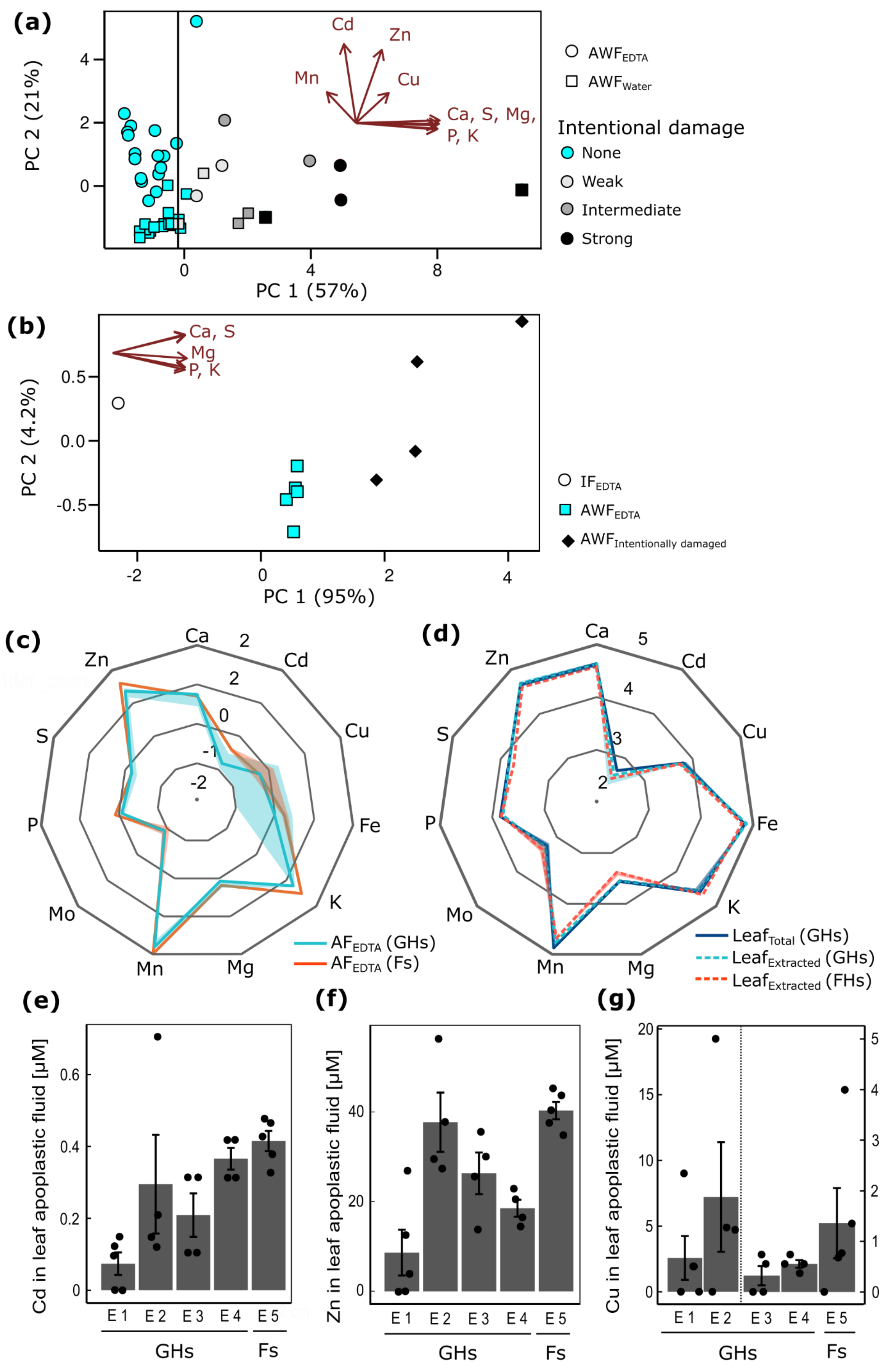

(a)
PC 2 (21%)
PC 1 (57%)
Cd
Zn
Mn
Cu
Ca, S, Mg, P, K
AWF_EDTA
AWF_Water
Intentional damage
None
Weak
Intermediate
Strong
(b)
PC 2 (4.2%)
PC 1 (95%)
Ca, S
Mg
P, K
IF_EDTA
AWF_EDTA
AWF_Intentionally damaged
(c)
Ca
Cd
Cu
Fe
K
Mg
Mn
Mo
P
S
Zn
AF_EDTA (GHs)
AF_EDTA (Fs)
(d)
Leaf_Total (GHs)
Leaf_Extracted (GHs)
Leaf_Extracted (FHs)
(e)
Cd in leaf apoplastic fluid [µM]
E 1
E 2
E 3
E 4
E 5
GHs
Fs
(f)
Zn in leaf apoplastic fluid [µM]
(g)
Cu in leaf apoplastic fluid [µM]

**Figure S4. Ionomes of leaf and leaf apoplast samples of *A. thaliana*. (a)** Effects of infiltration fluid type and of intentional leaf damage on the composition of apoplastic wash fluid. Shown is a PCA of the concentrations (*z*-scores of Log-transformed values) of nine elements in leaf apoplastic wash fluid (AWF), which identified two significant principal components, PC 1 (95%-confidence interval (CI): 50-63.6) and PC 2 (95%-CI: 18.8-26.8), with percentages of explained variation given in parentheses (bootstrapping with 999 replicates and 999 random permutations). Each datapoint represents one AWF sample obtained from the leaves of one or two plants, with several datapoints from each of four independent experiments (E 1 to E 4, see (e-g) below). PCA was conducted based on five macronutrient elements (Ca, K, Mg, P, and S with significant loadings on PC 1) and four trace metals (Cd and Zn with significant loadings on PC 2, as well as Cu and Mn). PC 1 separates AWF samples according to the degree of cytoplasmic contamination. PC 2 separates AWF samples obtained with water from those obtained with EDTA solution as the IF. The vertical line marks the threshold employed here to exclude samples potentially affected by cytoplasmic contamination from further analysis. **(b)** Test of the procedure developed in (a) on an independent set of AWF samples. PCA of the concentrations (*z*-scores of log-transformed values) of the five macronutrient elements chosen to detect cytoplasmic contamination in AWF (see (a)), for plants cultivated in a field-collected soil (E 5, see (e-g) below). The intensity of intentional damage was intermediate. Shown are the first two components PC 1 (significant) and PC 2, with percentages of explained variation given in parentheses (bootstrapping with 999 replicates, and 999 random permutations). All five elements have significant loadings on and significant correlations with PC 1 (*PCAtest* function, 999 bootstrap replicates, and 999 random permutations). **(c**, **d)** Ionomes of leaf apoplastic fluid (c), and of entire leaves before and after the extraction of apoplastic wash fluid (d). Shown (c) are median (line) and 25/75%iles (shadows) for each element as $Log_{10}$(concentration [mM] for Ca, S, P, Mg, and K; [µM] for Zn, Mo, Mn, Fe, Cu, and Cd; *n* = 17 samples for greenhouse soil (GHs, experiments E1-E4) and *n* = 4 samples for field-collected soil (Fs, experiment E 5, see e-g). Shown (d) are median (line) and 25/75%iles (shadows) for each element as $Log_{10}$(concentration [µg $g^{-1}$ DW] for Ca, S, P, Mg, and K; [ng $g^{-1}$ DW] for Zn, Mo, Mn, Fe, Cu, and Cd) in entire leaves (total): *n* = 6 samples from 2 independent experiments (GHs), leaves after AWF extraction (extracted): *n* = 28 samples from 4 independent experiments (GHs) and *n* = 5 samples from one experiment (Fs). (**e**, **f**, **g**) Concentrations of Cd (e), Zn (f), and Cu (g) in leaf AF of wild-type *A. thaliana*. Bars represent mean ± SD (*n* = 4 to 5 replicate pools of two plants), with one datapoint shown per replicate. Each bar is from an independent experiment (E 1 to E 5). *A. thaliana* (Col-0) were cultivated in short-day conditions for seven to eight weeks before the harvest of leaves. AF, leaf apoplastic fluid. IF, infiltration fluid (water or 100 µM EDTA).

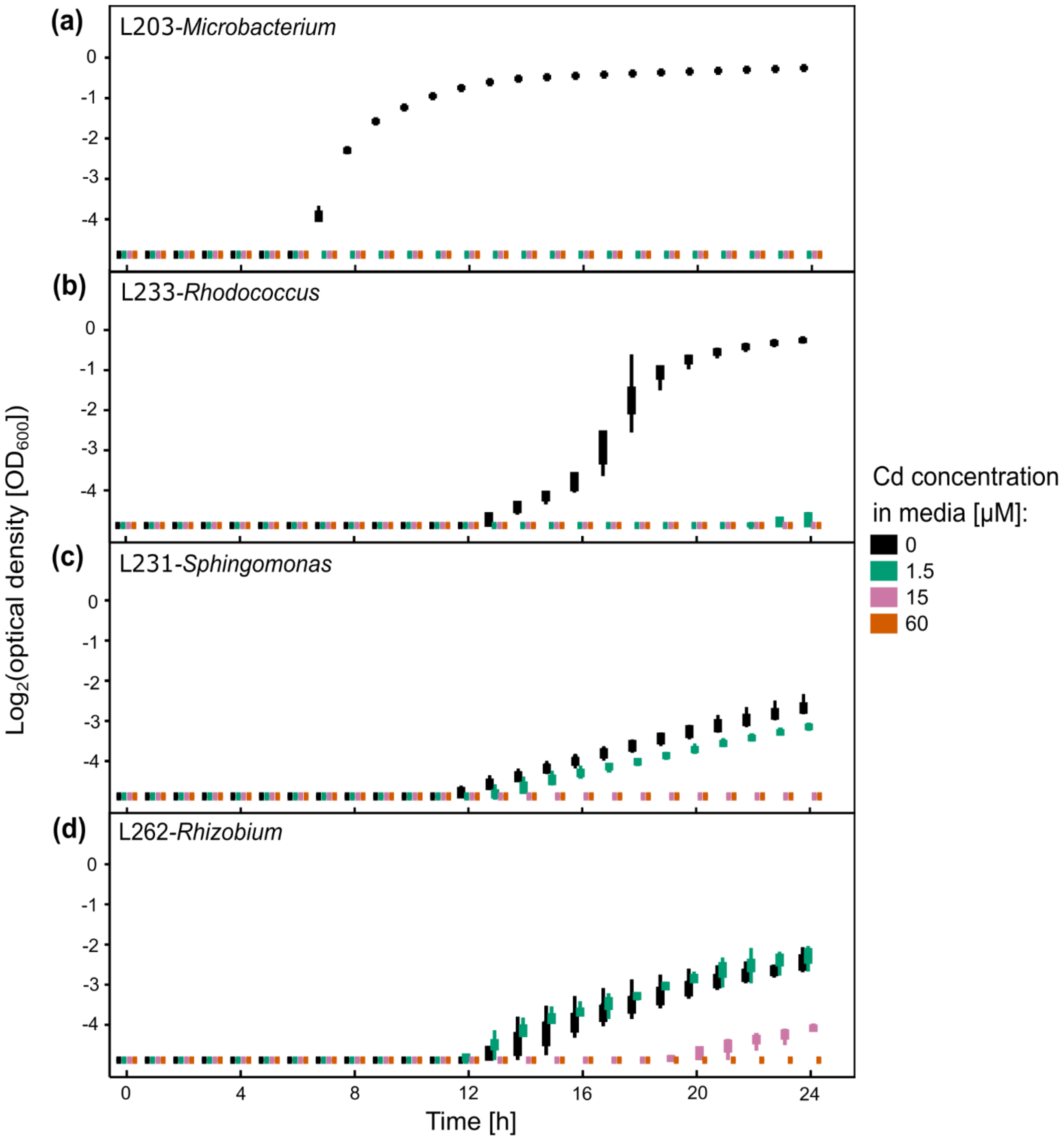


**Figure S5. Cd tolerance of keystone strains in liquid media. (a-d)** Shown are growth curves as time series of optical densities of single bacterial strains cultivated in liquid R2A media containing 0, 1.5, 15, or 60 µM Cd for the four keystone strains (a) L203-*Microbacterium*, (b) L233-*Rhodococcus,* (c) L231-*Sphingomonas*, and (d) L262-*Rhizobium* (boxplots of $n$ = 6 technical replicates). Data are from one experiment representative of a total of four independent experiments. The optical density was measured at 600 nm once per h during cultivation at 26°C for 24 h.

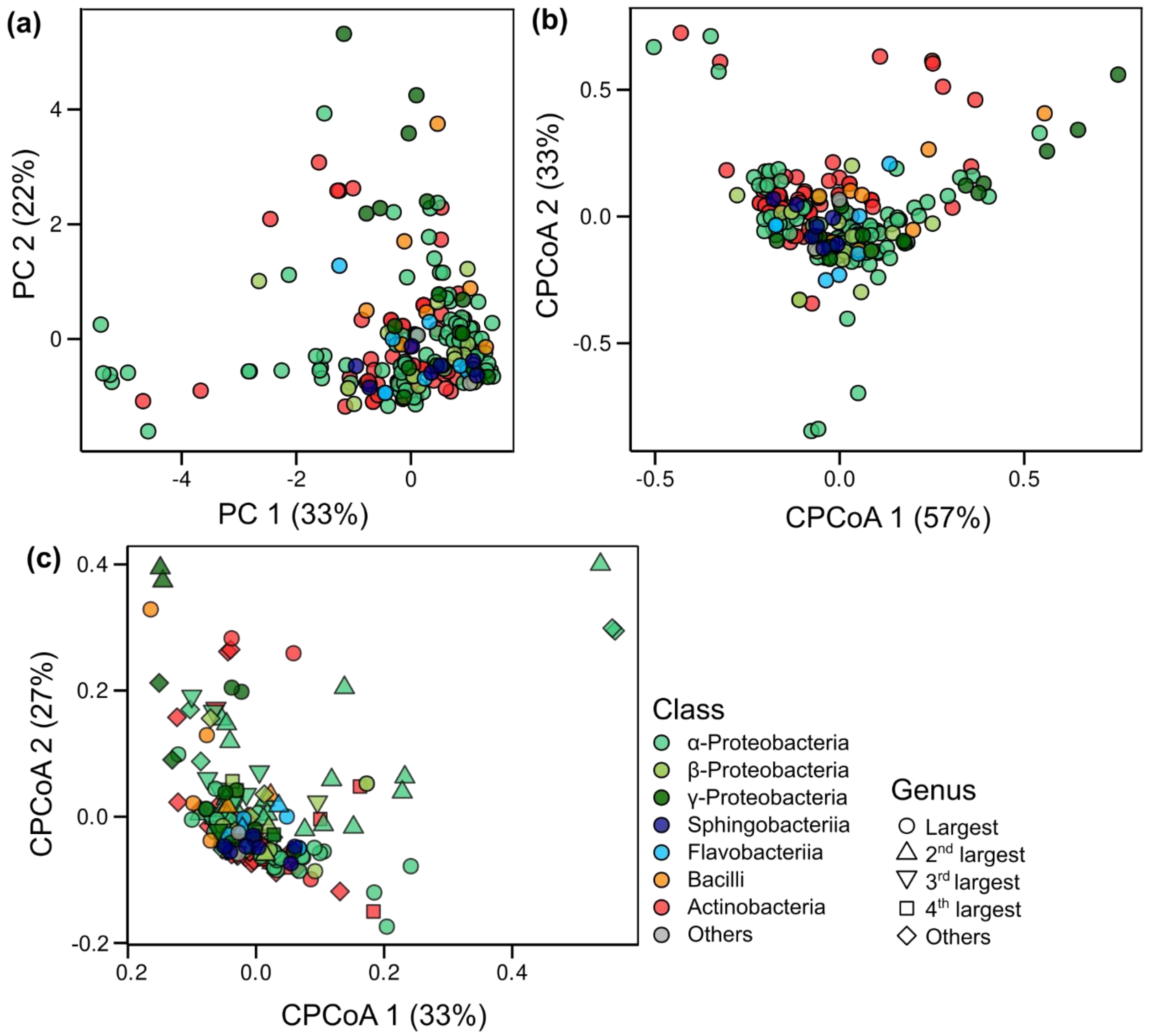


**Figure S6. Relationship between metal tolerance profiles and phylogeny of *At*-LSPHERE strains. (a)** PCA of median maximum permissive metal(loid) concentrations (MPC) of Cd, Zn, Cu, Mn, and arsenite, in the strains of the *At*-LSPHERE collection ($n$ = 3 independent experiments, Dataset S1). PC 1 was statistically significant, with significant loadings for MPC of Cu and Zn (bootstrapping with 999 replicates, and 999 random permutations). **(b**, **c)** Constrained principal coordinate analysis (CPCoA) of MPC (as in (a)), constrained by taxonomic class (B; 5.69% of the variance, $p = 0.03$), and by genus (C; 42.3% of the variance, $p < 0.001$). Statistically significant are CPCoA 1 ($p = 0.04$) in (b), and both CPCoA 1 ($p = 0.004$) and CPCoA 2 ($p = 0.01$) in (c), based on ANOVA-like permutation tests (999 permutations). Each datapoint (a-c) represents one strain, with coloring by phylogenetic class (Alphaproteobacteria, $n$ = 99; Betaproteobacteria, $n$ = 19; Gammaproteobacteria, $n$ = 16; Sphingobacteriia, $n$ = 7; Flavobacteriia, $n$ = 9; Deinococci, $n$ = 1; Bacilli, $n$ = 8; Actinobacteria, $n$ = 65). The shapes of datapoints in (c) are diversified by genus, reflecting genus size in the *At*-LSPHERE collection. The analyses were conducted on $z$-scores (a-c).

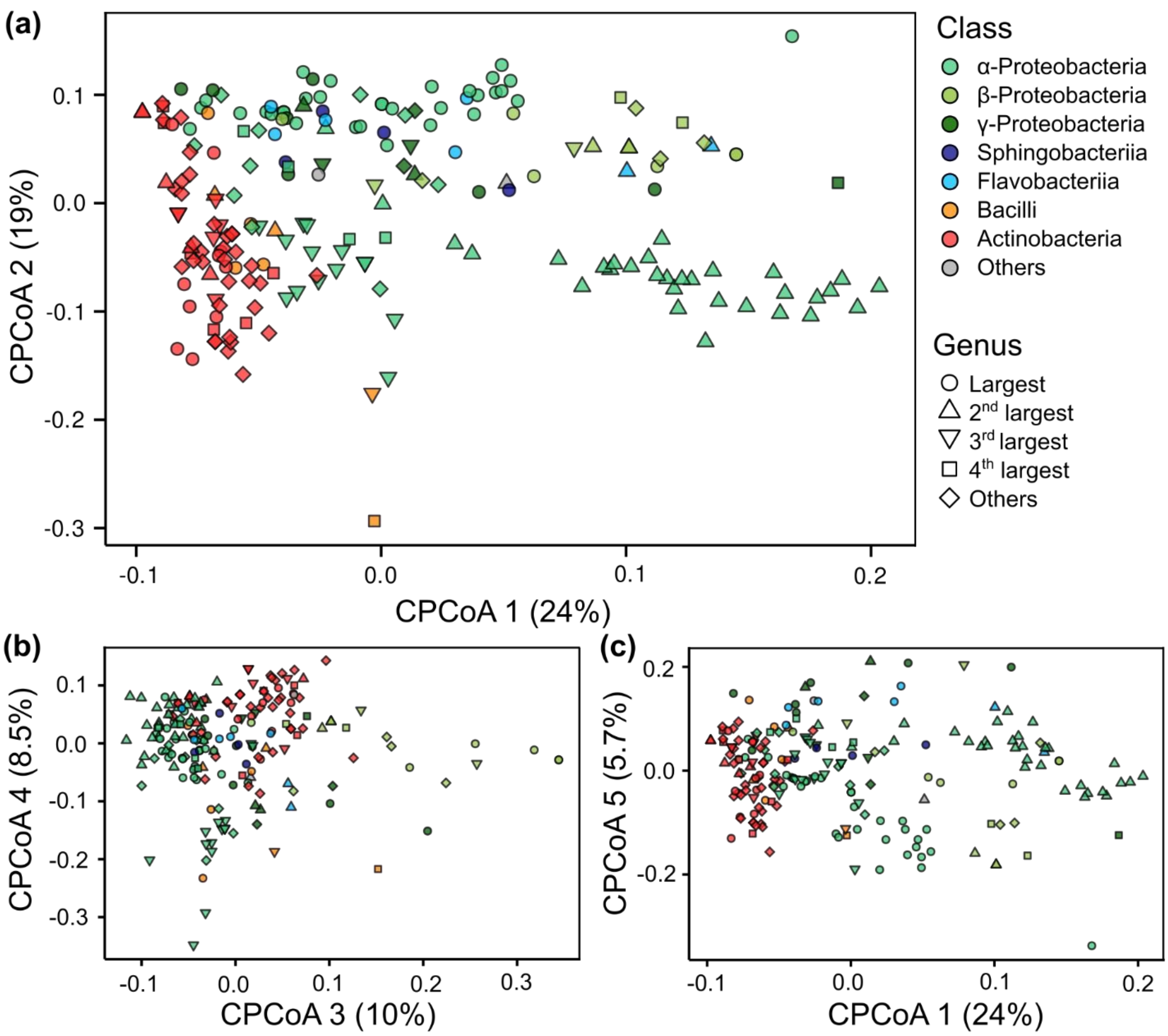


**Figure S7. Relationship between metal-related genes in the genome and phylogeny of *At*-LSPHERE strains. (a-c)** Constrained principal coordinate analysis (CPCoA), constrained by genus, with 1st and 2nd (a), 3rd and 4th (b), and 1st and 5th axes (c) shown ($p < 0.05$ in forward test, 999 permutations). An ANOVA-like permutation test for distance-based redundancy analysis was used to assess the significance of constraints (999 permutations, $p \leq 0.001$). Each datapoint represents one strain. The analysis was based on 100 metal-related KEGG Ortholog groups (KO), each of which was present in ≥ 8 bacterial strains (Datasets S2 and S3).

**Table S1. Composition of R2A medium.**

| **Compound** | **Company** | **Concentration [g $L^{-1}$ ]**[a] | **Concentration in low-P medium [g $L^{-1}$]**[a,b] |
|---|---|---|---|
| Casein acid hydrolysate | Merck KGaA, Darmstadt, Germany | 0.5 | 0.5 |
| Yeast extract | Becton, Dickinson and Company, United States, Franklin Lakes | 0.5 | 0.5 |
| Proteose peptone No.3 | Becton, Dickinson and Company, United States, Franklin Lakes | 0.5 | 0.5 |
| Glucose | Fisher Scientific GmbH, Schwerte, Germany | 0.5 | 0.5 |
| Starch | Merck KGaA, Darmstadt, Germany | 0.5 | 0.5 |
| di-Potassium hydrogen phosphate | Carl Roth GmbH + Co. KG, Karlsruhe, Germany | 0.3 | 0.03 |
| Sodium pyruvate | Sigma-Aldrich Chemie GmbH, Taufkirchen, Germany | 0.3 | 0.3 |
| HEPES | AppliChem GmbH, Darmstadt, Germany | - | 0.48 |
| Magnesium sulfate heptahydrate | Grüssing GmbH, Filsum, Germany | 0.05 | 0.05 |
| Bacto agar [c] | Becton, Dickinson and Company, United States, Franklin Lakes | 15 | 15 |
| Methanol (100%) | | 3.96 (5 mL) | 3.96 (5 mL) |

[a]pH adjusted to 7.2 with HCl; [b]low-phosphate medium used in arsenate concentration series (including also the associated 0 $As^{V}$ control); [c]omitted in liquid medium.

**Table S2. Metal salts used in tolerance screening on solid R2A medium.**

| **Metal salt** | **Company** | **Concentration [mM][a,c]** | **Concentration in low-P medium [mM][a,b,c]** |
|---|---|---|---|
| $CdSO_4$ | Life Technologies GmbH, Darmstadt, Germany | **0**, $1.5\times10^{-3}$, **0.015, 0.06, 0.12, 0.24, 0.48,** 1.5 | - |
| $CuSO_4$ | Sigma-Aldrich Chemie GmbH, Taufkirchen, Germany | **0,** 0.01, **0.1, 0.2, 0.4, 0,6, 1**, 2, 5, 10 | - |
| $ZnSO_4$ | VWR International GmbH, Darmstadt, Germany | **0,** 0.05, **0.1, 0.2, 0.4, 1, 2,** 10 | - |
| $Na_2HAsO_4$ | Merck KGaA, Darmstadt, Germany | - | **0,** 0.1, **1, 2,** 4, 5, **10, 20,** 50, 100 |
| $MnSO_4$ | RIEDEL-DE HAEN AG, Seelze-Hannover, Germany | **0,** 0.05, 0.25, 0.5, 1**, 2, 5, 10, 20 (or 50), 100** | - |
| $NaAsO_2$ | Merck KGaA, Darmstadt, Germany | **0,** 0.025, **0.25, 0.75, 1.5, 2,5, 5,** 10 | - |

[a]pH adjusted to 7.2 with HCl; [b]low-phosphate medium used in arsenate concentration series and its respective control; [c]$n \geq 3$ (**black and bold**), $n \leq 2$ (grey), $n$ = number of independent experiments.

| Table S3. Total concentrations of elements in R2A media as quantified by ICP-OES. | | | |
|---|---|---|---|
| **Elements** | **Concentration[a]** | **Conc. low-P medium[a,b]** | **Unit** |
| **Agar-solidified unamended control media[c]** | | | |
| Copper (Cu) | 0.33 ± 0.028 | 0.22 ± 0.11 | µM |
| Iron (Fe) | 8.1 ± 0.16 | 4.4 ± 1.0 | |
| Manganese (Mn) | 0.39 ± $7.9\times10^{-3}$ | 0.21 ± 0.049 | |
| Molybdenum (Mo) | 0.14 ± 0.013 | 0.088 ± $5.3\times10^{-3}$ | |
| Zinc (Zn) | 2.6 ± 0.29 | 1.3 ± 0.28 | |
| Cadmium (Cd) | 0.040 ± $2.5\times10^{-3}$ | 0.031 ± $1.6\times10^{-3}$ | |
| Calcium (Ca) | 0.75 ± 0.015 | 0.39 ± 0.097 | mM |
| Potassium (K) | 5.1 ± 0.14 | 1.7 ± 0.42 | |
| Magnesium (Mg) | 0.55 ± 0.011 | 0.39 ± 0.094 | |
| Phosphorus (P) | 2.3 ± 0.040 | 0.40 ± 0.10 | |
| Sulphur (S) | 4.9 ± 0.12 | 4.4 ± 1.1 | |
| **Liquid unamended control media[c]** | | | |
| Copper (Cu) | 0.049 ± $7.2\times10^{-3}$ | - | µM |
| Iron (Fe) | 1.0 ± 0.14 | - | |
| Manganese (Mn) | 0.020 ± $3.4\times10^{-4}$ | - | |
| Molybdenum (Mo) | 0.015 ± $2.0\times10^{-4}$ | - | |
| Zinc (Zn) | 1.3 ± 0.026 | - | |
| Cadmium (Cd) | $3.6\times10^{-4}$ | - | |
| Calcium (Ca) | 0.021 ± $1.5\times10^{-4}$ | - | mM |
| Potassium (K) | 4.1 ± 0.058 | - | |
| Magnesium (Mg) | 0.15 ± $2.1\times10^{-3}$ | - | |
| Phosphorus (P) | 2.3 | - | |
| Sulphur (S) | 0.59 ± 0.010 | - | |

[a]pH adjusted to 7.2 with HCl; [b]low-phosphate medium used for tolerance screening in arsenate concentration series and its respective control; [c]mean ± SD ($n$ = 3 independent samples)

| Table S4. Cd tolerances of individual strains of the synthetic communities. | |
|---|---|
| **Bacteria** | **Strain-wise median of maximum permissive Cd concentration [µM][a]** |
| **Cd-sensitive SynCom** | **15** |
| *L1-Plantibacter* | 15 |
| *L8-Frigoribacterium* | 15 |
| *L41-Pedobacter* | 15 |
| *L75-Bacillus* | 15 |
| *L108-Methylobacterium* | 15 |
| *L151-Microbacterium* | 15 |
| *L363-Brevundimonas* | 15 |
| *L399-Methylobacterium* | 15 |
| *L459-Methylophilus* | 15 |
| *L466-Methylobacterium* | 15 |
| **Intermediate SynCom** | **240** |
| *L16-Sphingomonas* | 240 |
| *L48-Pseudomonas* | 240 |
| *L78-Acidovorax* | 60 |
| *L87-Methylobacterium* | 240 |
| *L132-Pedobacter* | 240 |
| *L210-Agreia* | 240 |
| *L226-Sphingomonas* | 240 |
| *L278-Rhodococcus* | 240 |
| *L325-Leifsonia* | 180 |
| *L371-Rhizobium* | 240 |
| **Cd-tolerant SynCom** | **1,500** |
| *L10-Sphingomonas* | 480 |
| *L61-Duganella* | 480 |
| *L82-Flavobacterium* | 480 |
| *L90-Methylobacterium* | 1,500 |
| *L123-Methylobacterium* | 480 |
| *L137-Arthrobacter* | 1,500 |
| *L170-Pedobacter* | 480 |
| *L220-Variovorax* | 1,500 |
| *L337-Arthrobacter* | 1,500 |
| *L396-Bradyrhizobium* | 1,500 |

[a]Solid R2A media supplemented with 0, 1.5, 15, 60, 120, 240, 480, and 1,500 µM Cd; cultivation at RT for 20 d (*n* = 3).

**Table S5. The ionome of the *A. thaliana* leaf apoplastic fluid.**

| Experiment | Infiltration fluid (IF) | n[a] | Elements | Trace elements [µM] | | | | | | | Macronutrients [mM] | | | | |
|---|---|---|---|---|---|---|---|---|---|---|---|---|---|---|---|
| | | | | Cd | Cu | Fe | Mn | Mo | Pb | Zn | Ca | K | Mg | P | S |
| | | | Wavelength [nm] | 214.4 | 324.7 | 259.9 | 257.6 | 202 | 220.3 | 213.8 | 318.1 | 769.8 | 279 | 185.9 | 182 |
| GHs 1 | Water | 5 | **mean** | **n.d.** | **5.48** | **0.16** | **21.1** | **0.01** | **n.d.** | **8.55** | **6.98** | **22.4** | **1.18** | **0.58** | **0.78** |
| | | | SD | n.d. | 5.23 | 0.32 | 4.16 | 0.03 | n.d. | 2.71 | 1.27 | 3.91 | 0.18 | 0.14 | 0.29 |
| | EDTA | 5 | **mean** | **0.07** | **2.57** | **2.26** | **49.0** | **0.08** | **0.31** | **8.66** | **6.14** | **19.1** | **1.48** | **0.89** | **0.62** |
| | | | SD | 0.06 | 3.33 | 2.09 | 20.0 | 0.10 | 0.62 | 10.2 | 0.51 | 2.44 | 0.55 | 0.49 | 0.26 |
| GHs 2 | Water | 2 | **mean** | **n.d.** | **1.66** | **n.d.** | **41.5** | **0.06** | **0.10** | **6.95** | **7.32** | **23.5** | **1.65** | **0.93** | **0.82** |
| | | | SD | n.d. | 0.01 | n.d. | 11.7 | 0.06 | 0.10 | 1.46 | 2.42 | 2.66 | 0.54 | 0.15 | 0.09 |
| | EDTA | 4 | **mean** | **0.30** | **7.21** | **1.67** | **49.4** | **0.12** | **0.21** | **37.8** | **6.52** | **20.4** | **1.32** | **0.71** | **1.04** |
| | | | SD | 0.24 | 7.22 | 2.39 | 22.4 | 0.11 | 0.17 | 11.5 | 2.05 | 1.82 | 0.37 | 0.08 | 0.28 |
| GHs 3 | Water | 4 | **mean** | **n.d.** | **3.69** | **1.84** | **44.4** | **0.02** | **0.13** | **5.19** | **4.84** | **14.9** | **1.72** | **1.21** | **0.84** |
| | | | SD | n.d. | 6.28 | 2.32 | 7.06 | 0.01 | 0.17 | 0.94 | 1.22 | 2.88 | 0.42 | 0.44 | 0.35 |
| | EDTA | 4 | **mean** | **0.21** | **0.32** | **0.74** | **71.3** | **0.06** | **0.24** | **26.4** | **4.45** | **14.6** | **1.64** | **0.97** | **0.62** |
| | | | SD | 0.10 | 0.33 | 1.28 | 20.2 | 0.06 | 0.15 | 8.04 | 0.95 | 1.15 | 0.30 | 0.20 | 0.05 |
| GHs 4 | Water | 4 | **mean** | **n.d.** | **0.60** | **2.59** | **60.5** | **0.08** | **0.54** | **8.39** | **5.37** | **18.5** | **1.98** | **0.73** | **0.34** |
| | | | SD | n.d. | 0.72 | 0.88 | 20.8 | 0.12 | 0.36 | 4.38 | 1.20 | 2.88 | 0.42 | 0.04 | 0.04 |
| | EDTA | 4 | **mean** | **0.37** | **0.55** | **1.63** | **85.0** | **0.15** | **0.10** | **18.6** | **2.37** | **13.0** | **1.18** | **0.82** | **0.46** |
| | | | SD | 0.05 | 0.13 | 1.07 | 10.3 | 0.05 | 0.14 | 3.30 | 0.15 | 1.08 | 0.05 | 0.13 | 0.17 |
| Fs 1 | EDTA | 5 | **mean** | **0.40** | **1.18** | **1.89** | **91.3** | **0.12** | **0.08** | **39.8** | **5.52** | **30.9** | **1.47** | **1.29** | **0.66** |
| | | | SD | 0.06 | 1.43 | 0.74 | 8.48 | 0.04 | 0.01 | 4.73 | 0.89 | 2.00 | 0.10 | 0.26 | 0.11 |
| **Apoplastic Fluid Minimum** | | | | 0.07 | 0.32 | 0.16 | 21.1 | 0.0 | 0.1 | 5.2 | 2.4 | 13.0 | 1.2 | 0.6 | 0.3 |
| **Apoplastic Fluid Median** | | | | 0.30 | 1.2 | 1.7 | 71 | 0.12 | 0.21 | 26 | 5.5 | 22 | 1.6 | 0.9 | 0.78 |
| **Apoplastic Fluid Maximum** | | | | 0.40 | 7.2 | 2.6 | 91 | 0.15 | 0.31 | 40 | 7.3 | 31 | 2.0 | 1.3 | 0.84 |
| **$EC_5$ (curve fit-based)[b]** | | | | 4.2 | 110 | n.a. | 3,000 | n.a. | n.a. | 83 | n.a. | n.a. | n.a. | n.a. | n.a. |
| **Concentrations in solid R2A[c]** | | | | varied | varied | 8.1 | varied | 0.14 | n.a. | varied | 0.75 | 5.1 | 0.55 | 2.3 | 4.9 |

Plants were cultivated on our greenhouse soil (GHs) or a field soil mixture (Fs). [a]Number of replicates (leaves taken from a replicate set of 2-3 plants); [b]concentration at which 5% of the tested bacterial strains were unable to grow; [c]see Tables S2 and S3; n.d.: under the limit of reliable quantification; n.a.: not analyzed. The concentrations of As in AWF samples were below our detection limits and are thus omitted here.

**Table S6. Soil ionomes for the greenhouse soil and the field-collected soil mixtures used for plant cultivation.**

| Elements | Greenhouse soil (GHs, pH 5.8)[a] | | | | | | Field soil mixture (Fs, pH 5.9)[a] | | | | | |
|---|---|---|---|---|---|---|---|---|---|---|---|---|
| | Total | | Extractable | | Exchangeable | | Total | | Extractable | | Exchangeable | |
| | **Mean[b]** | **SD[b,d]** | **Mean[b]** | **SD[b,d]** | **Mean[b]** | **SD[b,d]** | **Mean[b]** | **SD[b,d]** | **Mean[b]** | **SD[b,d]** | **Mean[b]** | **SD[b,d]** |
| **Trace elements [mg kg$^{-1}$ DW[c]][b]** | | | | | | | | | | | | |
| **Aluminium (Al)** | **3,100** | 530 | **110** | 27 | **6.3** | 0.7 | **3,200** | 400 | **82** | 3.8 | **1.0** | 0.3 |
| **Boron (B)** | **2.5** | 0.3 | **40** | 1.2 | **0.6** | 0.1 | **2.6** | 0.2 | **39** | 0.1 | **0.3** | 0.0 |
| **Cadmium (Cd)** | **n.d.** | n.d. | **n.d.** | n.d. | **n.d.** | n.d. | **n.d.** | n.d. | **n.d.** | n.d. | **n.d.** | n.d. |
| **Copper (Cu)** | **6.7** | 0.9 | **0.7** | 0.4 | **0.1** | 0.0 | **3.8** | 0.4 | **0.8** | 0.0 | **0.0** | 0.0 |
| **Iron (Fe)** | **2,700** | 440 | **63** | 25 | **3.5** | 0.40 | **4,400** | 470 | **47** | 0.80 | **1.0** | 0.2 |
| **Manganese (Mn)** | **57** | 9.6 | **25** | 6.5 | **1.1** | 0.2 | **69** | 15 | **13** | 1.4 | **0.8** | 0.10 |
| **Nickel (Ni)** | **4.6** | 0.70 | **0.7** | 0.06 | **n.d.** | n.d. | **8.9** | 1.0 | **0.7** | 0.06 | **n.d.** | n.d. |
| **Zinc (Zn)** | **12** | 0.90 | **6.3** | 0.7 | **1.2** | 0.1 | **14** | 1.4 | **3.7** | 0.4 | **0.5** | 0.1 |
| **Macronutrients [mg kg$^{-1}$ DW[c]][b]** | | | | | | | | | | | | |
| **Calcium (Ca)** | **9,600** | 1,000 | **10,000** | 1,300 | **3,600** | 130 | **4,200** | 540 | **3,800** | 480 | **1,800** | 210 |
| **Potassium (K)** | **800** | 100 | **450** | 81 | **370** | 36 | **800** | 72 | **210** | 24 | **180** | 21 |
| **Magnesium (Mg)** | **770** | 99 | **390** | 71 | **180** | 11 | **1,500** | 130 | **180** | 31 | **110** | 14 |
| **Phosphorus (P)** | **550** | 65 | **400** | 67 | **290** | 38 | **300** | 18 | **190** | 16 | **90** | 16 |
| **Sulphur (S)** | **530** | 50 | **150** | 32 | **31** | 2.6 | **230** | 29 | **64** | 8.8 | **19** | 1.4 |

[a]Soil treated as in experiments; [b]$n$ = 5 replicate soil samples; [c]DW: dry mass; [d]SD: standard deviation; n.d.: below the limit of reliable quantification of 0.1 mg Cd kg$^{-1}$ or 0.25 mg Ni kg$^{-1}$.

**Table S7. Leaf ionomes before and after AWF extraction for *A. thaliana* grown in greenhouse soil or field soil mixture.**

| Elements | Greenhouse soil substrate (GHs) | | | | Field soil substrate (Fs) | |
|---|---|---|---|---|---|---|
| | Bulk leaves[a] | | After AWF extraction[b] | | After AWF extraction[c] | |
| | **Mean** | **SD[d]** | **Mean** | **SD[f]** | **Mean** | **SD[f]** |
| **Trace elements [µg g$^{-1}$ DW[e]]** | | | | | | |
| **Cadmium (Cd)** | **0.57** | 0.084 | **0.46** | 0.071 | **0.35** | 0.0028 |
| **Copper (Cu)** | **6.2** | 0.37 | **5.8** | 0.36 | **5.6** | 0.34 |
| **Iron (Fe)** | **77** | 8.5 | **83** | 5.2 | **62** | 10 |
| **Manganese (Mn)** | **82** | 4.3 | **66** | 6.6 | **49** | 12 |
| **Molybdenum (Mo)** | **2.0** | 0.027 | **2.0** | 0.15 | **2.3** | 0.50 |
| **Zinc (Zn)** | **50** | 1.5 | **46** | 5.7 | **40** | 2.1 |
| **Macronutrients [µg g$^{-1}$ DW[e]]** | | | | | | |
| **Calcium (Ca)** | **38,000** | 3,300 | **38,000** | 3,300 | **38,000** | 6,100 |
| **Potassium (K)** | **44,000** | 3,000 | **42,000** | 2,800 | **47,000** | 7,700 |
| **Magnesium (Mg)** | **3,300** | 350 | **3,300** | 140 | **2,400** | 440 |
| **Phosphorus (P)** | **6,900** | 270 | **6,700** | 450 | **6,600** | 300 |
| **Sulphur (S)** | **6,800** | 230 | **6,300** | 500 | **5,300** | 330 |

[a]Untouched leaves, calculated from $n$ = 3 pools of leaves, each from one to two replicate plants, experiment E4; [b]calculated from $n$ = 7 pools of leaves, each from one to two replicate plants, experiment E4; [c]calculated from $n$ = 5 pools of leaves, each from one to two replicate plants, experiment E5; [e]DW: dry biomass; [f]SD: standard deviation

**Table S8. Combined effects of cadmium and copper toxicity on keystone strains.**

| Cd µM | Cu µM | E1 | E2 | E3 |
|---|---|---|---|---|
| L203-*Microbacterium* | | | | |
| 0 | 1[a] | T: 0, V: 0 | T: 0, V: + | T: +, V: 0 |
| 0 | 10[a] | T: 0, V: 0 | T: 0, V: + | T: +, V: 0 |
| 0 | 100[a] | T: 0, V: - | T: 0, V: + | T: 0, V: - |
| **0.5** | **0[a]** | **T: -, V: -** | **T: -, V: -** | **T: -, V: -** |
| 0.5 | 1[b] | T: 0, V: 0 | n.e. | n.e. |
| 0.5 | 10[b] | T: 0, V: 0 | n.e. | n.e. |
| 0.5 | 100[b] | T: -, V: - | n.e. | n.e. |
| **2** | **0[a]** | **T: -, V: -** | **T: -, V: -** | **T: -, V: -** |
| 2 | 1[b] | T: 0, V: - | n.e. | n.e. |
| 2 | 10[b] | T: -, V: - | n.e. | n.e. |
| 2 | 100[b] | T: -, V: - | n.e. | n.e. |
| L231-*Sphingomonas* | | | | |
| 0 | 1[a] | T: 0, V: 0 | T: 0, V: 0 | T: 0, V: 0 |
| 0 | 10[a] | T: 0, V: 0 | T: 0, V: 0 | T: 0, V: 0 |
| 0 | 100[a] | T: 0, V: 0 | T: 0, V: 0 | T: 0, V: 0 |
| **0.5** | **0[a]** | **T: -, V: -** | **T: 0, V: -** | **T: -, V: 0** |
| 0.5 | 1[b] | T: 0, V: + | T: 0, V: + | T: +, V: 0 |
| 0.5 | 10[b] | T: +, V: + | T: 0, V: + | T: +, V: 0 |
| 0.5 | 100[b] | T: 0, V: + | T: 0, V: + | T: 0, V: 0 |
| **2** | **0[a]** | **T: -, V: -** | **T: -, V: -** | **T: -, V: -** |
| 2 | 1[b] | T: +, V: + | T: +, V: + | T: 0, V: 0 |
| 2 | 10[b] | T: 0, V: + | T: +, V: + | T: 0, V: 0 |
| 2 | 100[b] | T: +, V: + | T: +, V: + | T: 0, V: 0 |
| L233-*Rhodococcus* | | | | |
| 0 | 1[a] | T: 0, V: 0 | T: 0, V: 0 | T: 0, V: 0 |
| 0 | 10[a] | T: 0, V: 0 | T: 0, V: 0 | T: 0, V: 0 |
| 0 | 100[a] | T: 0, V: - | T: 0, V: - | T: 0, V: - |
| **0.5** | **0[a]** | **T: -, V: -** | **T: 0, V: 0** | **T: -, V: -** |
| 0.5 | 1[b] | T: 0, V: 0 | T: 0, V: 0 | T: 0, V: 0 |
| 0.5 | 10[b] | T: 0, V: 0 | T: 0, V: 0 | T: -, V: - |
| 0.5 | 100[b] | T: -, V: - | T: -, V: - | T: -, V: - |
| **2** | **0[a]** | **T: -, V: -** | **T: -, V: -** | **n.d.** |
| 2 | 1[b] | n.e. | n.e. | n.d. |
| 2 | 10[b] | n.e. | n.e. | n.d. |
| 2 | 100[b] | n.e. | n.e. | n.d. |
| L262-*Rhizobium* | | | | |
| 0 | 1[a] | T: 0, V: 0 | T: 0, V: 0 | T: 0, V: 0 |
| 0 | 10[a] | T: 0, V: 0 | T: 0, V: 0 | T: 0, V: 0 |
| 0 | 100[a] | T: 0, V: - | T: 0, V: - | T: -, V: - |
| **0.5** | **0[a]** | **T: 0, V: 0** | **T: 0, V: 0** | **T: 0, V: 0** |
| 0.5 | 1[b] | T: 0, V: 0 | T: 0, V: 0 | T: 0, V: 0 |
| 0.5 | 10[b] | T: 0, V: 0 | T: 0, V: 0 | T: 0, V: 0 |
| 0.5 | 100[b] | T: -, V: - | T: -, V: - | T: -, V: - |
| **2** | **0[a]** | **T: 0, V: 0** | **T: 0, V: 0** | **n.d.** |
| 2 | 1[b] | T: 0, V: 0 | T: 0, V: 0 | n.d. |
| 2 | 10[b] | T: 0, V: 0 | T: 0, V: 0 | n.d. |
| 2 | 100[b] | T: -, V: - | T: -, V: - | n.d. |

+/0/-: Bacterial growth promoted/unchanged/inhibited. [a]Comparison to the metal-free control. [b]Comparison to the respective Cu-free condition with the same Cd concentration (highlighted in **bold**). T: Time at which $OD_{600}$ reached a threshold (6-fold the SD of all measurements during the initial 4 h of cultivation, with significance testing by one-way ANOVA with Tukey's multiple comparisons *post hoc* test). V: Visually detected change in trajectory or shape of growth curve relative to untreated controls. n.d.: No data; n.e.: No evaluation possible

**Table S9. Combined effects of cadmium, zinc and manganese on keystone strains.**

| Cadmium *x* µM | Metals added *x*/50/10 µM Cd/Zn/Mn | E1 | E2 | E3 | E4 |
|---|---|---|---|---|---|
| | | L203-*Microbacterium* | | | |
| 0 | Mn[a] | T: 0; V: 0 | T: +; V: + | T: 0; V: 0 | n.d. |
| **0** | **Zn[a]** | **T: -; V: -** | **T: 0; V: 0** | **T: 0; V: -** | n.d. |
| 0 | Zn+Mn[b] | T: 0; V: 0 | T: -; V: - | T: -; V: - | n.d. |
| **0.5** | **Cd[a]** | **T: -; V: -** | **T: -; V: -** | **T: -; V: -** | n.d. |
| **0.5** | **Cd+Zn[c]** | **T: 0; V: 0** | **T: +; V: +** | **T: +; V: +** | n.d. |
| 0.5 | Cd+Mn[c] | T: +; V: + | T: ++; V: ++ | T: +; V: + | n.d. |
| 0.5 | Cd+Zn+Mn[d] | T: +; V: + | T: 0; V: 0 | T: +; V: + | n.d. |
| **5** | **Cd[a]** | **n.d.** | **T: -; V: -** | **T: -; V: -** | **T: -; V: -** |
| **5** | **Cd+Zn[c]** | **n.e.** | **T: +; V: +** | **T: 0; V: 0** | **T: 0; V: 0** |
| 5 | Cd+Mn[c] | n.e. | T: +; V: + | T: 0; V: 0 | T: 0; V: 0 |
| 5 | Cd+Zn+Mn[d] | n.e. | T: +; V: + | T: 0; V: 0 | T: 0; V: 0 |
| **20** | **Cd[a]** | **T: -; V: -** | **T: -; V: -** | **T: -; V: -** | n.d. |
| **20** | **Cd+Zn[c]** | **n.e.** | **n.e.** | **n.e.** | n.d. |
| 20 | Cd+Mn[c] | n.e. | n.e. | n.e. | n.d. |
| 20 | Cd+Zn+Mn[d] | n.e. | n.e. | n.e. | n.d. |
| | | L231-*Sphingomonas* | | | |
| 0 | Mn[a] | T: 0; V: 0 | T: 0; V: + | T: 0; V: 0 | n.d. |
| **0** | **Zn[a]** | **T: -; V: -** | **T: 0; V: 0** | **T: 0; V: -** | n.d. |
| 0 | Zn+Mn[b] | T: -; V: - | T: 0; V: 0 | T: 0; V: 0 | n.d. |
| **0.5** | **Cd[a]** | **T: -; V: -** | **T: -; V: -** | **T: -; V: -** | **T: -; V: -** |
| **0.5** | **Cd+Zn[c]** | **T: -; V: -** | **T: 0; V: -** | **T: 0; V: 0** | **T: -; V: -** |
| 0.5 | Cd+Mn[c] | T: 0; V: 0 | T: 0; V: 0 | T: +; V: + | T: 0; V: 0 |
| 0.5 | Cd+Zn+Mn[d] | T: -; V: - | T: 0; V: 0 | T: +; V: + | T: 0; V: 0 |
| **5** | **Cd[a]** | **n.d.** | **T: -; V: -** | **T: -; V: -** | **T: -; V: -** |
| **5** | **Cd+Zn[c]** | **n.e.** | **n.e.** | **T: -; V: -** | n.d. |
| 5 | Cd+Mn[c] | n.e. | n.e. | T: -; V: - | n.d. |
| 5 | Cd+Zn+Mn[d] | n.e. | n.e. | T: -; V: - | n.d. |
| **20** | **Cd[a]** | **T: -; V: -** | **T: -; V: -** | **T: -; V: -** | n.d. |
| **20** | **Cd+Zn[c]** | **n.e.** | **n.e.** | **n.e.** | n.d. |
| 20 | Cd+Mn[c] | n.e. | n.e. | n.e. | n.d. |
| 20 | Cd+Zn+Mn[d] | n.e. | n.e. | n.e. | n.d. |
| | | L233-*Rhodococcus* | | | |
| 0 | Mn[a] | T: +; V: + | T: 0; V: 0 | T: 0; V: 0 | n.d. |
| **0** | **Zn[a]** | **T: -; V: 0** | **T: -; V: -** | **T: 0; V: -** | n.d. |
| 0 | Zn+Mn[b] | T: 0; V: 0 | T: 0; V: 0 | T: 0; V: 0 | n.d. |
| **0.5** | **Cd[a]** | **T: -; V: -** | **T: -; V: -** | **T: -; V: -** | n.d. |
| **0.5** | **Cd+Zn[c]** | **T: 0; V: +** | **T: 0; V: 0** | **n.e.** | n.d. |
| 0.5 | Cd+Mn[c] | T: +; V: + | T: 0; V: + | n.e. | n.d. |
| 0.5 | Cd+Zn+Mn[d] | T: +; V: + | T: 0; V: + | T: 0; V: + | n.d. |
| **5** | **Cd[a]** | **T: -; V: -** | **n.d.** | **T: -; V: -** | **T: -; V: -** |
| **5** | **Cd+Zn[c]** | **n.e.** | **n.e.** | **n.e.** | n.d. |
| 5 | Cd+Mn[c] | n.e. | n.e. | n.e. | n.d. |
| 5 | Cd+Zn+Mn[d] | n.e. | n.e. | n.e. | n.d. |
| **20** | **Cd[a]** | **T: -; V: -** | **T: -; V: -** | **T: -; V: -** | n.d. |
| **20** | **Cd+Zn[c]** | **n.e.** | **n.e.** | **n.e.** | n.d. |
| 20 | Cd+Mn[c] | n.e. | n.e. | n.e. | n.d. |
| 20 | Cd+Zn+Mn[d] | n.e. | n.e. | n.e. | n.d. |
| | | L262-*Rhizobium* | | | |
| 0 | Mn[a] | T: +; V: + | T: 0; V: 0 | T: 0; V: 0 | |
| **0** | **Zn[a]** | **T: -; V: -** | **T: -; V: -** | **T: 0; V: -** | n.d. |
| 0 | Zn+Mn[b] | T: +; V: + | T: 0; V: 0 | T: +; V: ++ | n.d. |

| | | | | | |
|---|---|---|---|---|---|
| **0.5** | **Cd[a]** | **T: -; V: -** | **T: 0; V: 0** | **T: -; V: -** | n.d. |
| **0.5** | **Cd+Zn[c]** | **T: -; V: -** | **T: -; V: -** | **T: 0; V: 0** | n.d. |
| 0.5 | Cd+Mn[c] | T: +; V: + | T: 0; V: 0 | T: +; V: + | n.d. |
| 0.5 | Cd+Zn+Mn[d] | T: +; V: + | T: +; V: + | T: +; V: + | n.d. |
| **5** | **Cd[a]** | **T: 0; V: 0** | **T: -; V: -** | **T: 0; V: -** | n.d. |
| **5** | **Cd+Zn[c]** | **T: -; V: -** | **T: 0; V: -** | **T: 0; V: -** | n.d. |
| 5 | Cd+Mn[c] | T: 0; V: 0 | T: +; V: + | T: 0; V: + | n.d. |
| 5 | Cd+Zn+Mn[d] | T: +; V: + | T: +; V: + | T: 0; V: + | n.d. |
| **20** | **Cd[a]** | **T: -; V: -** | **T: -; V: -** | **T: -; V: -** | n.d. |
| **20** | **Cd+Zn[c]** | **T: -; V: -** | **T: -; V: -** | **T: 0; V: 0** | n.d. |
| 20 | Cd+Mn[c] | n.d. | n.d. | n.d. | n.d. |
| 20 | Cd+Zn+Mn[d] | T: +; V: + | T: +; V: + | T: +; V: + | n.d. |

+/0/-: Bacterial growth promoted/unchanged/inhibited. [a]Comparison to control medium. [b]Comparison to Zn alone (highlighted in **bold**); [c]Comparison to the same concentration of Cd alone (highlighted in **bold** and underlined); [d]Comparison to combination of Cd and Zn at the same concentrations alone (highlighted in ***bold*** and in italics); T: Time at which $OD_{600}$ reached a threshold (6-fold the SD of all measurements during the initial 4 h of cultivation, with significance testing by one-way ANOVA with Tukey's multiple comparisons *post hoc* test). V: Visually detected change in trajectory or shape of growth curve relative to untreated controls. n.d.: No data; n.e.: No evaluation possible.

**Table S10. Known metal-related genes associated with metal tolerances in the *At*-LSPHERE collection.**

| Short name[a] | Function | $Cd^{II}$ | $Cu^{II}$ | $Zn^{II}$ | $As^{III}$ | $Mn^{II}$ | $As^{V}$ |
|---|---|---|---|---|---|---|---|
| *ABC.ZM.A* | ABC $Zn^{2+}$/$Mn^{2+}$ uptake, ATP-binding | n.s. | n.s. | n.s. | n.s. | + | n.s. |
| *ABC.ZM.P* | … permease | n.s. | n.s. | n.s. | n.s. | + | n.s. |
| *ABC.ZM.S* | … substrate-binding | n.s. | n.s. | n.s. | n.s. | + | n.s. |
| *arnF* | L-Ara4N lipid A mod. export (IM) | n.s. | - | n.s. | n.s. | n.s. | n.s. |
| *arnT..pmrK* | Lipid A L-Ara4N transferase | n.s. | n.s. | n.s. | n.s. | + | n.s. |
| *arsA..ASNA1..GET3* | ATPase for ArsB-$As^{III}$ efflux | -- | n.s. | n.s. | n.s. | n.s. | n.s. |
| *arsC1..arsC* | Arsenate reductase | + | - | - | n.s. | n.s. | n.s. |
| *arsH* | NADPH-FMN $As^{III}$ oxidase prod. $As^{V}$ | + | n.s. | n.s. | n.s. | n.s. | ++ |
| *ATOX1..ATX1..copZ..golB* | Cu-chaperone for $Cu^{+}$ efflux | n.s. | - | - | n.s. | + | n.s. |
| *baeS..smeS* | Sensor histidine kinase | n.s. | n.s. | n.s. | n.s. | + | n.s. |
| *bccA..pccA* | Biotin-/propionyl-CoA carboxylase | - | n.s. | n.s. | n.s. | n.s. | n.s. |
| *comB* | DNA uptake (natural competence) | -- | + | n.s. | n.s. | n.s. | n.s. |
| *comC* | Competence-stimulating | n.s. | n.s. | - | n.s. | + | n.s. |
| *copA..ATP7* | P-type $Cu^{+}$-exporting ATPase | n.s. | n.s. | n.s. | n.s. | + | n.s. |
| *copC..pcoC* | Periplasmic Cu-binding chaperone | n.s. | + | + | n.s. | n.s. | n.s. |
| *csoR..ricR* | Cu-sensing transcriptional repressor | - | n.s. | n.s. | n.s. | n.s. | n.s. |
| *cueR* | Cu-responsive activator of copA/cueO | n.s. | n.s. | n.s. | n.s. | + | n.s. |
| *cusB..silB* | Periplasmic adaptor for Cu/Ag efflux | + | n.s. | n.s. | n.s. | n.s. | n.s. |
| *cutA* | Cu binding/tolerance | n.s. | n.s. | n.s. | n.s. | n.s. | + |
| *czcA* | CzcCBA (Co/Zn/Cd) efflux system inner membrane transporter | + | n.s. | n.s. | n.s. | n.s. | n.s. |
| *czcB* | … periplasmic membrane fusion prot. | + | n.s. | n.s. | n.s. | n.s. | n.s. |
| *czcC* | … outer membrane channel | + | n.s. | n.s. | n.s. | n.s. | n.s. |
| *czcD..zitB* | $Zn^{2+}$ exporter (low-level Zn resistance) | n.s. | n.s. | n.s. | n.s. | n.s. | ++ |
| *DPYS..dht..hydA* | Incl. $H_2$-producing FeFe-hydrogenase | n.s. | n.s. | - | n.s. | n.s. | n.s. |
| *mdtA* | RND-type multidrug efflux pump periplasmic adaptor | + | n.s. | - | n.s. | + | n.s. |
| *mdtB* | … transporting component | + | n.s. | - | n.s. | n.s. | n.s. |
| *mdtC* | … transporting component | + | n.s. | - | n.s. | n.s. | n.s. |
| *mdtD* | MFS polyamine/small mol. exporter | n.s. | - | n.s. | n.s. | + | + |
| *mdtG* | MFS antibiotic efflux pump | n.s. | n.s. | n.s. | n.s. | + | n.s. |
| *mdtH* | MFS transporter; fluoroquinolone res. | n.s. | n.s. | n.s. | n.s. | n.s. | -- |
| *mdtP* | MATE $Na^{+}$-driven multidrug exporter | n.s. | n.s. | n.s. | n.s. | - | n.s. |
| *mmcO* | Periplasmic MCO detoxifying $Cu^{+}$ | n.s. | n.s. | n.s. | n.s. | n.s. | ++ |
| *mntR* | Transcriptional regulator of Mn uptake | n.s. | - | - | n.s. | + | + |
| *oprM..emhC..ttgC..cusC..adeK..smeF..mtrE..cmeC..gesC* | RND efflux system outer membrane channel | n.s. | n.s. | n.s. | n.s. | - | n.s. |
| *PCCA..pccA* | See *bccA..pccA* | n.s. | n.s. | - | n.s. | n.s. | + |
| *pcoB..copB* | Outer membrane Cu export protein | n.s. | n.s. | n.s. | n.s. | - | n.s. |
| *pcoD* | IM Cu transporter periplasmic detox. | - | + | n.s. | n.s. | n.s. | n.s. |
| *PTS-Mtl-EIIA..mtlA..cmtB* | PTS: mannitol phosphorylation | -- | n.s. | + | n.s. | n.s. | n.s. |
| *PTS-Mtl-EIIB..mtlA....cmtA* | … EIIB domain | n.s. | n.s. | + | n.s. | n.s. | n.s. |
| *PTS-Mtl-EII..mtlA..cmtA* | … EIIABC/ mannitol transp./enzyme | -- | n.s. | + | n.s. | n.s. | n.s. |
| *smtB* | Zn-responsive transcriptional repressor | -- | n.s. | n.s. | n.s. | n.s. | n.s. |
| *sitA* | Periplasmic $Mn^{2+}$/$Fe^{2+}$-binding | n.s. | n.s. | n.s. | n.s. | + | n.s. |
| *sitB* | … ATP-binding | n.s. | n.s. | n.s. | n.s. | + | n.s. |
| *sitC* | … $Mn^{2+}$/$Fe^{2+}$ permease | n.s. | n.s. | n.s. | n.s. | + | n.s. |
| *sitD* | … permease or stability factor | n.s. | n.s. | n.s. | n.s. | + | n.s. |
| *tolC* | OM channel for RND/MFS/ABC/TolC-mediated efflux | + | n.s. | n.s. | n.s. | n.s. | n.s. |
| *zntA* | $Zn^{2+}$/$Pb^{2+}$/$Cd^{2+}$-exporting ATPase | + | n.s. | n.s. | n.s. | n.s. | n.s. |
| *zntB* | $Zn^{2+}$ uptake/export (CDF) | n.s. | - | - | n.s. | n.s. | n.s. |
| *znuA* | High-affinity periplasmic $Zn^{2+}$-binding | - | n.s. | n.s. | n.s. | n.s. | n.s. |
| *znuC* | ZnuABC permease/ATPase | n.s. | - | - | n.s. | n.s. | n.s. |
| *zur* | Zn-sensor controlling *znu* genes | + | n.s. | n.s. | n.s. | n.s. | ++ |

[a]Short names of genes, for which median MPC of positive strains differed from median MPC of negative strains lacking an orthologue (≥ 2-fold, $p_{adj} < 0.05$) for ≥ 1 metal(loid). n.s.: no significant effect; **+**: 2- to 10-fold increase in MPC; **++:** ≥ 10-fold increase in MPC; **-**: 2- to 10-fold decrease in MPC; **--:** ≥ 10-fold decrease in MPC; .., separates short names linked to identical KEGG Ortholog id. IM/OM: inner/outer membrane.

### Supporting Information - Supplementary Methods Text

#### Plant cultivation

*Arabidopsis thaliana* pre-cultivation was conducted in 10 cm x 10 cm x 11 cm square pots (~ 100 seeds per pot) of standard soil treated once with Previcur Energy (Bayer CropScience, Monheim, Germany). After transfer to individual pots, plants were fertilized with 0.3% (v/v) Wuxal Super 8-8-6 (Hauert MANNA Düngewerk GmbH, Nürnberg, Germany) every 14 d and watered with a suspension of *Bacillus thuringiensis israelensis* (Neudomück, W. Neudorff GmbH KG, Emmerthal, Germany) once a week.

#### Additional metal(loid) tolerance testing

For growth tests of SynCom strains and SynComs on solid media, 5-µL aliquots of each stock were spotted manually, followed by incubation in darkness at RT for 10 d.

#### Isolation of apoplastic wash fluid from leaves of *A. thaliana*

All handling of plant tissues was conducted using plastic tweezers. For each sample, leaves of ≥ 75% full-grown length that appeared healthy and unscathed were cut off at the petiole using a razor blade, pooled from two *A. thaliana* plants, rinsed in ultrapure water, blotted dry, and weighed. For vacuum infiltration, leaves were immersed in infiltration fluid (IF) inside a 1-L side arm vacuum flask containing 1 L of IF and covered by a plastic mesh cover that was weighed down by a glass rod placed on top of the mesh. Subsequently, after recovering the leaves and blotting them dry gently with soft tissue paper, five to eight leaves of one sample were placed next to one another on a 10 cm x 10 cm segment of Parafilm ("M" strips; PECHINEY PLASTIC PACKAGING, Chicago, United States of America), rolled gently around the outer surface of 5-mL pipette tips, and the arrangements were then placed tip-down into 50-mL polypropylene tubes. Apoplastic wash fluid (AWF) was extracted by centrifuging, followed by measuring the volume of each obtained AWF sample and weighing the remaining extracted leaves to identify possible differences in hydration status between plants and experiments. The recovered AWF was transferred into 1.5-mL tubes, centrifuged at 15,000x*g* for 5 min, and the supernatant transferred into a fresh 1.5-mL tube and stored at -80°C. Cytoplasmic contamination was assessed by quantifying glucose-6-phosphate dehydrogenase activity (MAK014-1KT, Sigma-Aldrich, Taufkirchen, Germany) according to the manufacturer's instructions (Dataset S5), and later using ionomic profiles. To identify the effects of cytoplasmic contamination on AWF ionomic profiles, we intentionally damaged leaf samples (in duplicate) using both IF solutions in each experiment. For weak simulated damage, we released the vacuum faster. For intermediate simulated damage, we additionally handled the leaves harshly after infiltration. For strong simulated damage, we additionally pressed the leaves against the 5-mL pipette tips. In each experiment, IF was sampled from the vacuum flasks following the removal of the leaves after infiltration to quantify baseline contamination originating from the leaf surfaces, and the materials and solutions used. Compared to a PCA of ionomic profiles of AWF (Fig. S4; see Methods below), we found that G6PDH activity was a far less sensitive marker for the detection of cytoplasmic contamination (Dataset S5).

**Multi-element analyses**

For multi-element analysis, aliquots (100 to 300 µL) of AWF were dried in 1.5-mL polypropylene tubes at 60°C for 7 d, and subsequently acid-digested by adding 500 µL 65% (w/w) $HNO_3$ and incubating RT overnight, then heating to 60°C for 1 h, 80°C for 1 h and 100°C for 1 h, with automated mixing at 300 rpm for 30 s every 5 min (Thermomixer Comfort, Eppendorf SE, Hamburg, Germany). Dried leaves (in paper bags, 60°C for 7 d, followed by RT > 3 d) were homogenized with 3 to 5 acid-washed ceramic beads (1.4 mm diameter) in 1.5-mL polypropylene tubes at 30 $s^{-1}$ for 30 s (Retsch MM300, RETSCH GmbH, Haan, Germany). Aliquots of approximately 20 mg dried leaf powder were weighed into perfluoroalkoxy alkane (PFA) vessels and digested in 3 mL 65% (w/w) $HNO_3$ using microwave-assisted digestion (1600 W, ramp time 20 min, hold time 15 min (190°C)) (MARSX$_{\text{PRESS}}$, CEM Corporation, Matthews, USA). After cooling to RT in ambient air, all digests were transferred into 15-mL polypropylene tubes and filled up to a final volume of 5 mL with ultrapure water. Fifty-mL aliquots of liquid R2A medium were dried in PFA vessels at 100°C and subsequently digested as described for leaves. Aliquots of around 120 mg agar were weighed into 15-mL polypropylene tubes, incubated in 2 mL 65% (w/w) $HNO_3$ at RT overnight and filled up to a total volume of 10 mL with ultrapure water. For the quantification of element concentrations by ICP-OES, calibrations were done using a blank and series of five multielement standards that were manually pipetted from single-element standard solutions (AAS Standards; Bernd Kraft, Duisburg, Germany) for 13 (AWF from experiments on greenhouse soil) or 16 (AWF from experiments on the natural soil mix) elements commonly detected in Arabidopsis and expected to be present in the apoplast, and 17 elements for analyses of bulk leaves. The precision of measurements was validated by measuring sample blanks and intermediate calibration standards, as well as for the analysis of leaves additionally a certified reference material (Virginia tobacco leaves, INCT-PVTL 6; Institute of Nuclear Chemistry and Technology, Warsaw, Poland) and an *A. thaliana* lab-internal reference material. Soil total and HCl-extractable fractions were prepared and analyzed as described (Stein *et al.*, 2017).

#### Details of data analysis and visualization

Depletion or enrichment of taxonomic classes was detected using Fisher's exact test (obtaining *q*-values using the *qvalue* package). For combined metal treatments, growth curves were fitted with generalized additive models with integrated smoothness estimation. The `cor.test()` function was used to test for a correlation between strain-wise Cd tolerance (MPC) and the counts of Cd tolerance-related gene and gene copies obtained using `mutate()` from *tidyverse*. Gene presence/absence calls were organized using the `pivot_longer()` function from the *tidyr* package. With a modified `GRdrawDRC()` function and the *ggplot2* package, data were plotted alongside dose-response curves. To show the metal tolerances of the keystone strains, spider charts were generated using Microsoft Excel. The heatmap plot was generated with the `heatmap.2()` function from the *gplots* package, with a customized dendrogram created with the *colorspace* (for color palette) and the *dendextend* packages. The correlation, PCA and CPCoA results of were plotted using the `ggbiplot()` and the `ggplot()` functions from the packages *ggbiplot* and *ggplot2*. For each strain and metal(loid), the MPC was normalized to the $EC_{50}$ of the full *At*-LSPHERE collection (see Table 1), linked to the tree, and plotted using the *ggtree* and *ggfruit* packages.
